# A Comprehensive Genetic Toolkit for Rapid and High-Resolution Engineering of Metabolic Pathways in *Zymomonas mobilis*

**DOI:** 10.1101/2025.10.29.685439

**Authors:** Melissa Poma, Matteo Marchetti, Cathal Burns, Andreas Zannetou, Emma M. Riley, Frank Sargent, Ulugbek Azimov, Jose L. Munoz-Munoz, Ciarán L. Kelly

## Abstract

The limited availability of reliable genetic tools has constrained the precise engineering of *Zymomonas mobilis*, a promising bacterium for the sustainable bioproduction of fuels and commodity chemicals. Here, we present a comprehensive, modular genetic toolkit that enables high-resolution, combinatorial control of gene expression in *Z. mobilis*. We constructed and characterised a synthetic promoter library spanning the broadest dynamic range reported for this organism, alongside a suite of strong transcriptional terminators that effectively insulate genetic cassettes and prevent transcriptional readthrough. To achieve predictable translation across diverse genetic contexts, we further implemented bicistronic design (BCD) elements, allowing context-independent control of expression. Promoter activities were shown to be stable under both aerobic and anaerobic conditions, highlighting the robustness of the regulatory parts across physiologically relevant environments. The toolkit was integrated into the Start-Stop Assembly system, facilitating high-throughput construction of multi-gene pathways. We demonstrate its utility through the rapid assembly and testing of the possible expression space of genes encoding a 2,3-butanediol biosynthetic pathway. Collectively, this genetic toolkit now allows the high-resolution engineering of multi-enzyme pathways in *Z. mobilis* for industrial biotechnology applications.

## Introduction

*Zymomonas mobilis* is a facultatively anaerobic, non-pathogenic, Gram-negative bacterium characterised by an exceptional ethanol production that reaches up to 98% of the theoretical maximum yield ^1^. This ability is attributed to the rapid consumption of glucose via the Entner-Doudoroff (ED) pathway ^2^, a less common and less energy-efficient route in Gram-negative bacteria compared to the Embden-Meyerhof (EMP) pathway, which yields two ATPs per glucose contrary to one from the ED pathway ^3^. The carbon and electron flux that is not invested into additional ATP production is instead directed towards ethanol fermentation, resulting in higher ethanol yields ^4^, which outperform those of EMP-equipped organisms, including the main ethanologenic competitor *Saccharomyces cerevisiae* ^5^.

A few studies have tackled the metabolic engineering of *Z. mobilis* for the conversion of abundant, low-cost biomass feedstocks into ethanol and other value-added chemicals ^6–11^. Early engineering efforts successfully expanded the natural substrate range of *Z. mobilis* to xylose ^12–16^ and arabinose ^17,18^, the two most abundant sugars in hemicellulose ^19^.

Additional work has focused on the heterologous expression of metabolic pathways for the production of various commodity chemicals, including 2,3-butanediol ^20^, poly-3- hydroxybutyrate ^21^, acetoin ^22^, lactate ^6,23^ and isobutanol ^24^.

Although the engineering of metabolic pathways has been widely attempted in *Z. mobilis*, a limited number of regulatory elements have been described in the literature. The regulation of transcription has mostly been limited to a narrow set of inducible promoters (P*tet*, P*tac* and P*lac*) ^6,24–30^, and constitutive promoters native to the *Z. mobilis* glycolytic pathway (P*gap*, P*eno* and P*pdc*) ^6,11,18,20–22,24,31–33^. Thirty-eight native promoters and four synthetic promoters have recently been characterised ^23,26^; however, these showed a limited range of strengths, which may be insufficient for finely-tuned gene expression. No transcriptional terminators have been characterised in *Z. mobilis* to date and their use rarely reported ^23^. Finally, various Ribosome Binding Sites (RBSs) have been characterised in *Z. mobilis* ^23,26^, but, as these are known to be context-dependent ^34^, no assumption of translation rate can be made when used in the 5’UTR or other sequences.

In this study, we explored and expanded the transcriptional and translational space of *Z. mobilis* by developing a tailor-made molecular toolbox comprised of twenty-nine novel synthetic promoters, strong transcriptional terminators and a bicistronic context-independent RBS system. Due to the industrial relevance of fermentation in *Z. mobilis*, we compared a subset of promoters in aerobic and anaerobic conditions. We subsequently used the newly developed regulatory parts to explore the transcriptional design space of a heterologous pathway for the industrially-important precursor chemical 2,3-butanediol.

## Results

### A Synthetic Promoter Library (SPL) with a wide range of expression strengths

Libraries of synthetic promoters can be created rapidly and can explore expression levels otherwise unattainable from simply screening native promoters ^21^. To date, various promoter engineering techniques have been developed, including random and targeted nucleotide deletions or substitutions in a native promoter ^22^, full promoter randomisation ^23,24^, and randomisation of nucleotides surrounding the -35 and -10 consensus regions ^25–28^. We chose the latter technique as it has been shown to generate a large distribution of strengths in different microbial hosts ^25–28^. The -35 and -10 consensus regions of σ⁷⁰ promoters were predicted from the genome of *Z. mobilis* using the software BPROM ^35^, and the frequency of each nucleotide calculated through a custom Python script (Figure S1 in the supplemental information). Nucleotides with a frequency threshold above 60% were fixed in the -35 and -10 box, and where this threshold was not met, the next most common nucleotide was included until this threshold was reached. Where no conservation was observed, nucleotides were randomised to introduce sequence diversity.

Having established the core promoter regions, we randomised the surrounding nucleotides: 10 bp upstream of the -35 box, the 17 bp inter operator space, and 6 bp downstream of the - 10 box (Figure 1), for a total length of 45 bp. These conserved and randomised sequences were introduced upstream of a gene encoding sfGFP on a *Z. mobilis* expression plasmid (pMP005). At this stage, the nucleotide composition of each putative promoter sequence was unknown.

**Figure 1.**
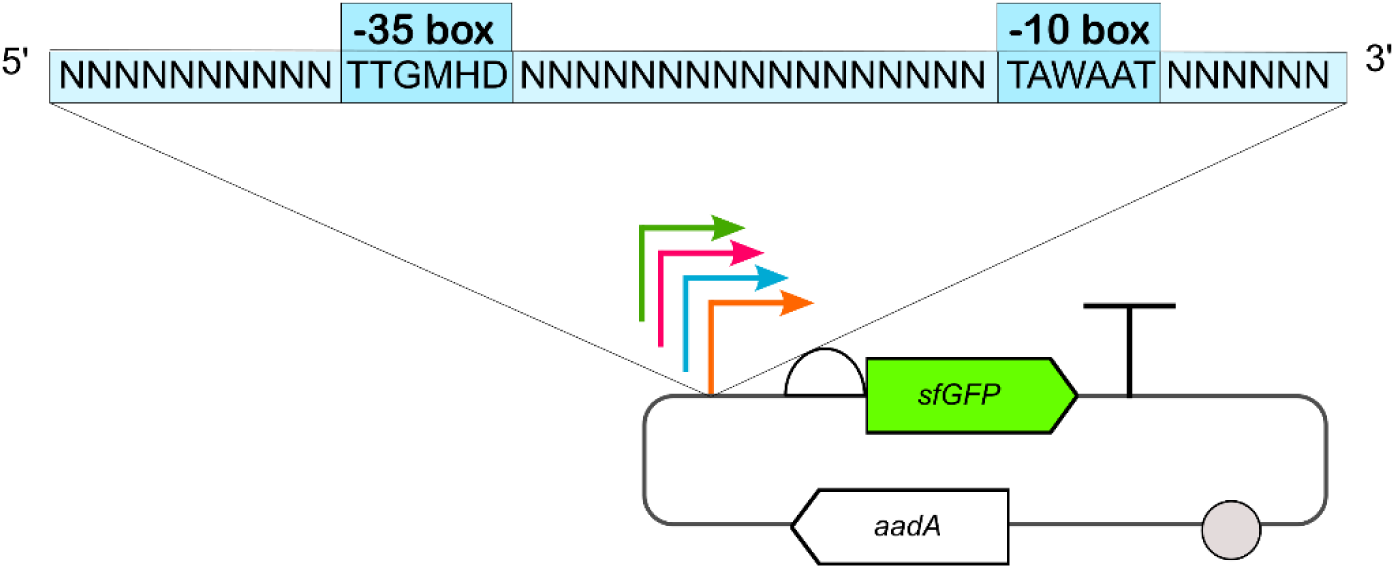
Synthetic promoter library (SPL) design. The most conserved nucleotides of the *Z. mobilis* -35 and – 10 boxes were fixed for the synthesis of partially-randomised synthetic promoters. ‘N’ = any nucleotide; ‘M’ = adenine or cytosine; ‘H’ = adenine, cytosine or thymine; ‘D’ = adenine, guanine or thymine; ‘W’ = adenine or thymine.

To compare the transcriptional strength of the synthetic promoters, negative and positive controls were designed and constructed. The plasmid pMP007, lacking a promoter and the gene encoding sfGFP, was used as negative control. Positive controls included two previously used constitutive promoters placed upstream of *sfGFP*: the T7A1 promoter (pMP005) ^36^ and the *glyceraldehyde-3-phosphate dehydrogenase* promoter (P*gap*) (pMP003), reported as the strongest native promoter and one of the most used for *Z. mobilis* engineering to date ^26^. Since P*gap* was predicted to be within a 305 bp genomic region ^26^, we generated a shorter 39 bp version containing just the predicted operator sites to reduce any effect that extra nucleotides may have on gene expression. This shorter putative sequence was renamed P*gap_*short, while the full 305 bp region was referred to as P*gap_*long.

The library was introduced into *Z. mobilis* by conjugation, using *E. coli* WM6026 as donor strain, and the fluorescence of eighty-four individual transconjugant colonies screened. This showed a wide range of fluorescence, spanning from values similar to the negative control, to values surpassing both positive controls (Figure S2 in the supplemental information). Twenty-nine individual colonies that showed >2% difference in fluorescence were taken forward for further characterisation by flow-cytometry in both *Z. mobilis* and *E. coli* WM6026 (Figure S3 in the supplemental information). This final screening confirmed that the newly generated SPL exhibited a broad significantly different range of transcriptional strengths in *Z. mobilis* (Figure 2) (Welch’s ANOVA on log₁₀-transformed data: F(32, 23.3) = 9699.6, p < 2.2 × 10⁻¹⁶). Post-hoc Welch pairwise tests with Benjamini–Hochberg correction identified multiple statistically distinct expression groups. Promoter activities spanned roughly 500-fold after back-transformation, with pMP045 and pMP034 yielding the highest mean expression (∼9 × 10³ a.u.) and pMP053 near background. Groupings are summarised by compact letter displays (Table S1 in supplementary information) and illustrated as ranked geometric means on a log₁₀ scale (Figure S4). Interestingly, the vast majority of the synthetic promoters had higher expression strength than the strongest native promoter described to date in other *Z. mobilis* genetic toolkits, P*gap*_long. P*gap*_short showed lower activity than the longer sequence.

**Figure 2.**
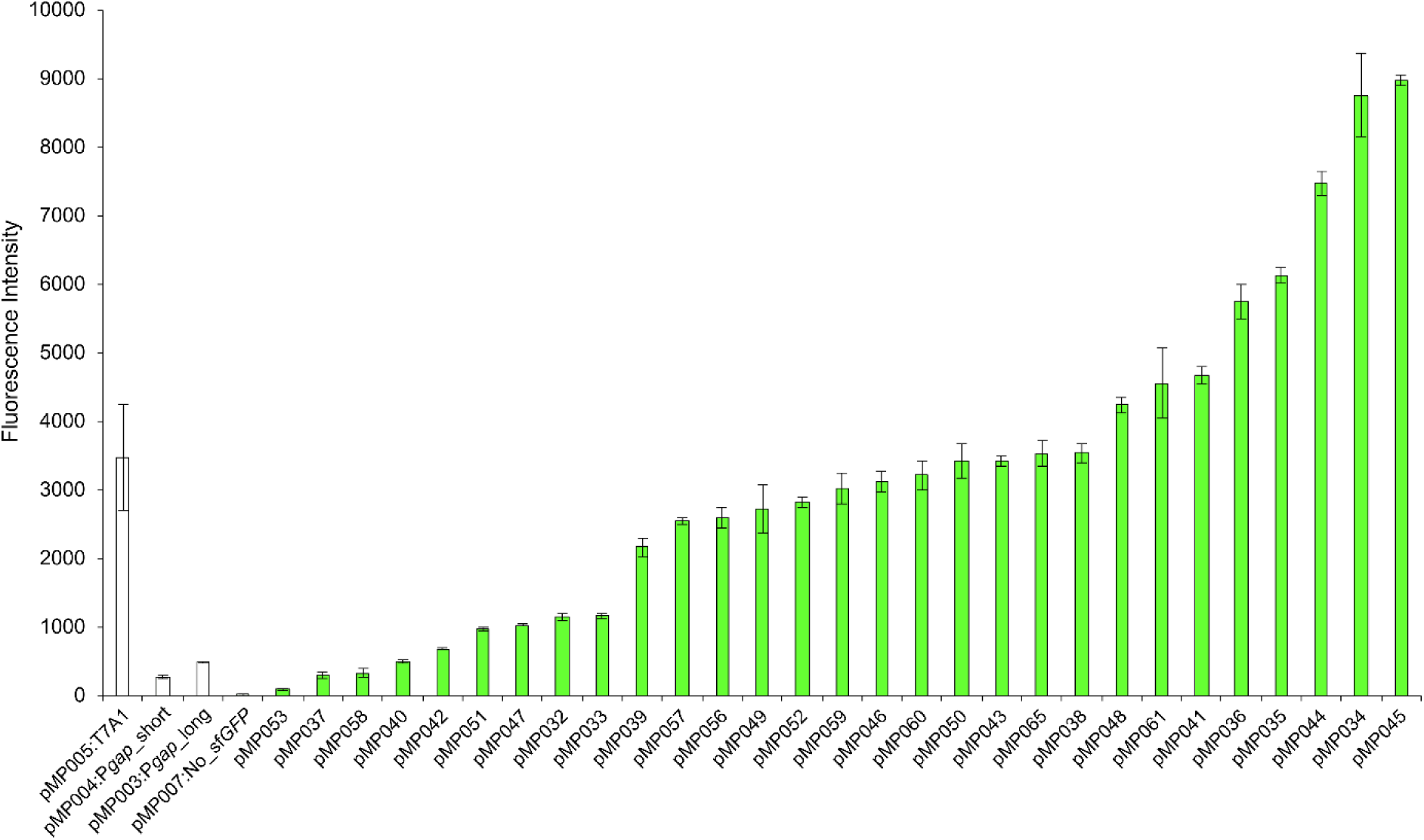
Fluorescence distribution of the SPL in *Z. mobilis*. Twenty-nine synthetic promoters were screened by flow-cytometry along with the following controls: pMP005:T7A1, pMP003:P*gap_*long, pMP004:P*gap_*short and pMP007:No_*sfGFP*. *Z. mobilis* recombinant strains were grown aerobically in ZRMG rich medium, at 30°C, in 96-well plates, shaking at 800 rpm. Samples were harvested during late exponential phase. Error bars represent the standard deviation of three independent biological replicates.

Unexpectedly, during Sanger sequencing of the final set of the synthetic promoter library (SPL) plasmids, it was noticed that the order of the –35 and –10 operator sequences had been unintentionally swapped during cloning (Figure S5 in the supplemental information), yet this atypical design still resulted in a large library of functional promoters with a wide range of transcriptional strengths. Additionally, it was noticed that some sequences were shorter than the expected 45 bp, but despite lacking nucleotides, they retained promoter activity and were therefore included for characterisation alongside the full-length sequences (Table S2 in the supplemental information).

To determine whether their strengths would differ if the canonical order was restored. To this end, we chose a weak (P18), medium (P19) and strong (P103) promoter and swapped their operator sequences. The modified promoters were renamed P18b, P19b and P103b, cloned upstream of *sfGFP*, and tested in *Z. mobilis* by flow-cytometry (Figure S6 in the supplemental information). P18b now exhibited the highest fluorescence; P103b was now in the middle of the three promoters; and P19b had the lowest promoter strength; demonstrating that although changing the order of the -35 and -10 boxes does alter promoter strength, both the standard and non-standard order of operators still allow transcription.

It has been suggested that reuse of promoters designed and characterised in *E. coli* should be sufficient for use in other Gram-negative bacteria. To test this, the fluorescence values of the SPL plasmids in *E. coli* strain WM6026 was compared to results in *Z. mobilis*. Fluorescence values in each organism were transformed to a percentage of the highest value observed in each organism and plotted on the *x*- and *y*- axes of a scatterplot. Concordance was assessed relative to a 1:1 line (*x = y*), which represents perfect agreement between the datasets. The fit of the datasets to the 1:1 line was calculated through the coefficient of determination (*R*^2^), which was equal to 0.05, The two groups had a coefficient of determination R2 to the 1:1 line of 0.05, suggesting deviation from concordance of this relationship. Additionally, as most of the datapoints fell above the 1:1 line, the SPL generally showed higher strength in *Z. mobilis* rather than E. coli. (Figure 3 in the supplemental information).

**Figure 3.**
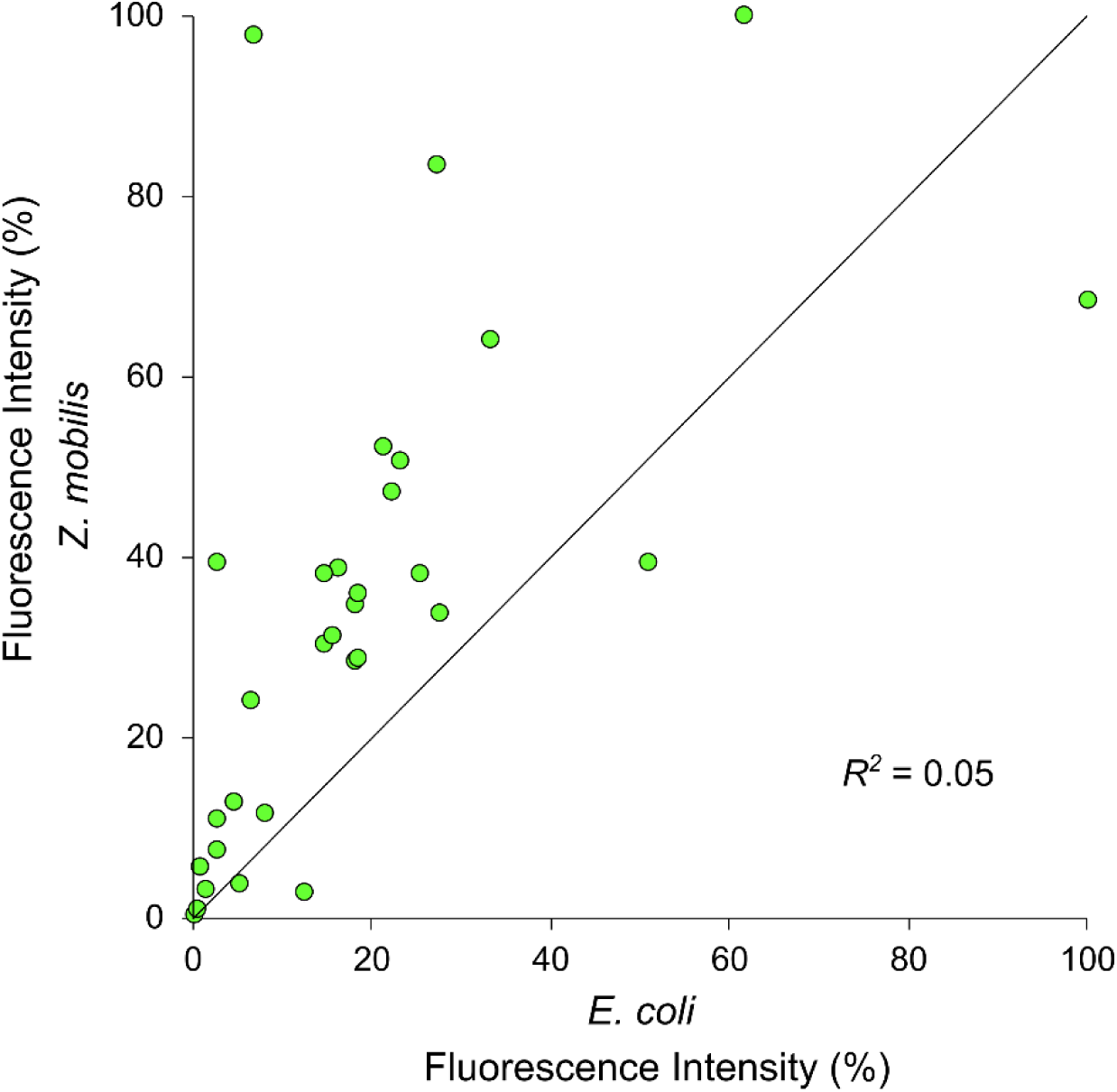
Concordance of synthetic promoter strengths between *Z. mobilis* and *E. coli*. The 1:1 line represents where the datapoints would fall if they were equivalent in the two organisms. The values were expressed in percentage in relation to the strongest promoters (100%) in *Z. mobilis* and *E. coli*.

### Anaerobic conditions do not affect the transcriptional strength of a subset of the SPL

As the production of reduced chemicals at high yields during fermentation is one of the key attractions to using *Z. mobilis* in biotechnology, we set out to investigate whether promoters characterised in aerobic conditions performed differently in anaerobic conditions. To this end, three promoters of the SPL, the weak P12, the medium P19 and the strong P103, were prepared for anaerobic screening by replacing the gene encoding the oxygen-dependent sfGFP with one encoding Y-FAST, an oxygen-independent protein that fluoresces upon binding of a fluorogen ^37^, resulting in plasmids pMP071, pMP072, pMP073 respectively. *Z. mobilis* transconjugants containing each of these plasmids were cultured without shaking in anaerobic conditions, and fluorescence measured by flow cytometry. Encouragingly, the pattern of strengths observed in aerobic conditions was maintained in anaerobiosis (Figure 4A), with GFP and Y-FAST values exhibiting a high concordance to the 1:1 line (*R*^2^ = 0.95), when expressed as a percentage of maximal fluorescence (Figure 4B).

**Figure 4.**
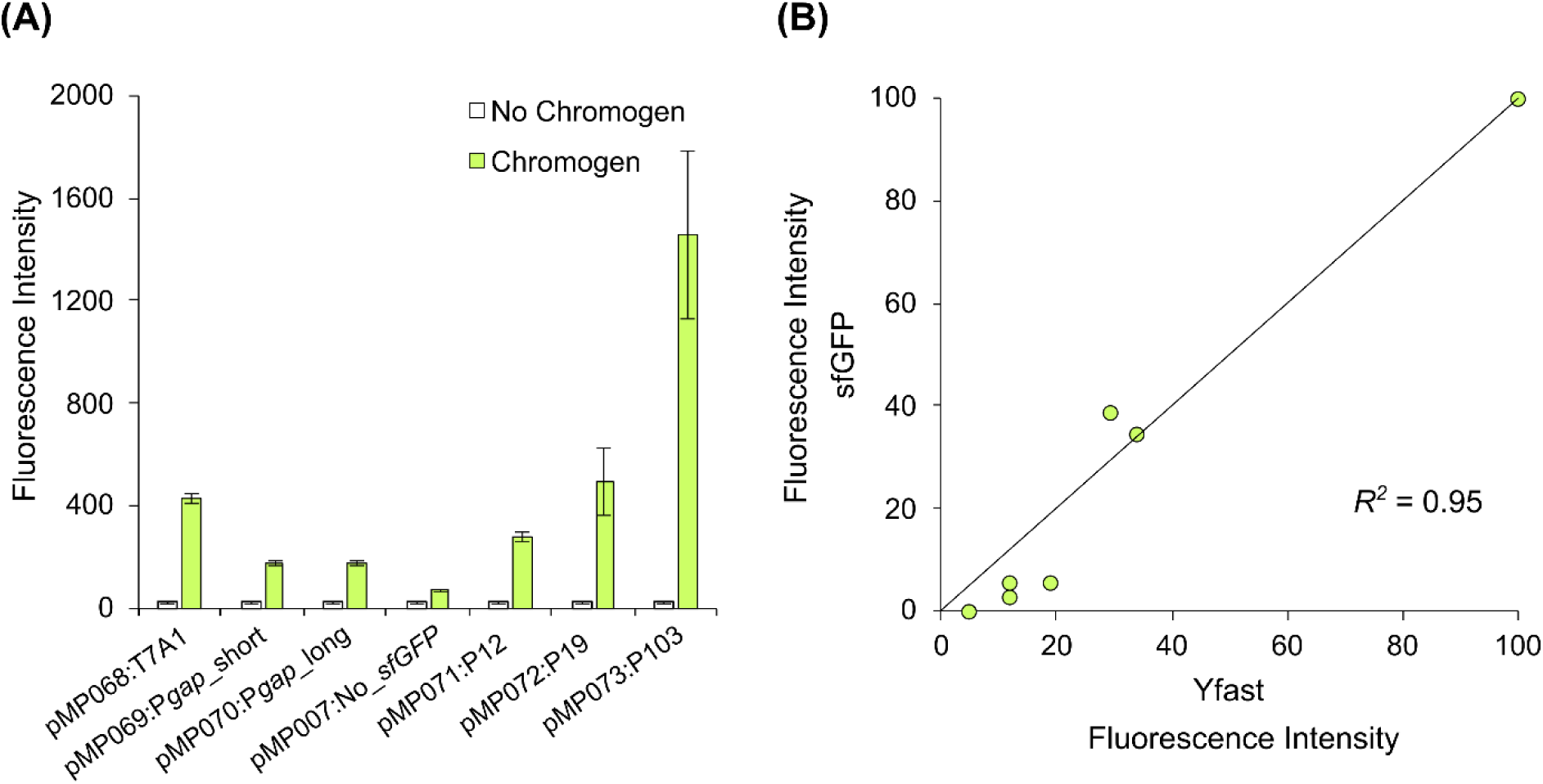
(A) Anaerobic screening of synthetic promoters in *Z. mobilis*. A weak (P12), a medium (P19) and a strong promoter (P103) from the aerobically tested SPL were cloned upstream of the *Y-FAST* reporter, generating the plasmids pMP071:P12:*Y-FAST*, pMP072:P19:*Y-FAST* and pMP073:P103:*Y-FAST*. These were tested along with the controls pMP68:T7A1:*Y-FAST*, pMP069:P*gap*_short:*Y-FAST*, pMP070:P*gap*_long:*Y-FAST* and pMP007:No*sfGFP*. Triplicates of recombinant *Z. mobilis* cultures were grown in ZRMG rich medium at 30 °C, in a 96-well plate in static mode to avoid oxygenation. Samples were harvested during late exponential phase and screened using flow-cytometry. Error bars represent the standard deviation of three independent biological replicates. **(B) Comparison of the synthetic promoter strengths in aerobiosis and anaerobiosis.** The fluorescence values of sfGFP and Y-FAST under the regulation of the T7A1, P*gap*_short, P*gap*_long, P12, P19 and P103 promoters were expressed in percentage in relation to the strongest promoter, P103 (100%). The sfGFP and Y-FAST values were respectively plotted on the *y-* and *x-* axis of a scatterplot with a 1:1 line.

### A collection of strong transcriptional terminators

No transcriptional terminators have been characterised to date in *Z. mobilis*, yet strong insulation between transcriptional units is required for the exploration of independent expression strengths and thus protein levels in multi-enzyme metabolic pathways. To address this gap, we sought to predict native Rho-independent (intrinsic) terminators and characterise them alongside heterologous and synthetic terminators previously described as strong in *E. coli* ^38^. Native intrinsic terminators were predicted using Softberry’s FindTerm software ^39^, and to minimise false positive predictions, open reading frames (ORFs) were removed from the input genome sequence. The secondary structures and Gibb’s free energy (Δ*G*) of putative native terminator sequences were analysed via UNAFold ^40^ and the ten sequences with the lowest Δ*G* were selected for phenotypic testing, along with two synthetic terminators and seven *E. coli* terminators (Table S3 in the supplemental information) ^38^. Since three of the ten putative terminators could not be cloned, the remaining seven were selected for thorough characterisation.

To characterise this selection of terminators, the dual-reporter plasmid pMM001 was built containing the genes encoding sfGFP and OpmCherry, under the regulation of the constitutive T7A1 promoter and two independent RBSs. Terminators were subsequently placed between the two reporters and the ratio of sfGFP/OpmCherry fluorescence used to assess transcriptional termination (Figure 5A). Plasmids containing *sfGFP* only (pMM005:*sfGFP*), *OpmCherry* only (pMM02:*OpmCherry*), and both fluorescent genes without intermediate terminators (pMM001:*sfGFP*:*OpmCherry*) were used as references. The fluorescence of *Z. mobilis* transconjugants was then studied using a microplate reader (Figure 5B). Heterologous or synthetic terminators blocked transcription with generally high efficiency, however six of the seven putative *Z. mobilis* terminators failed to reduce OpmCherry production.

**Figure 5.**
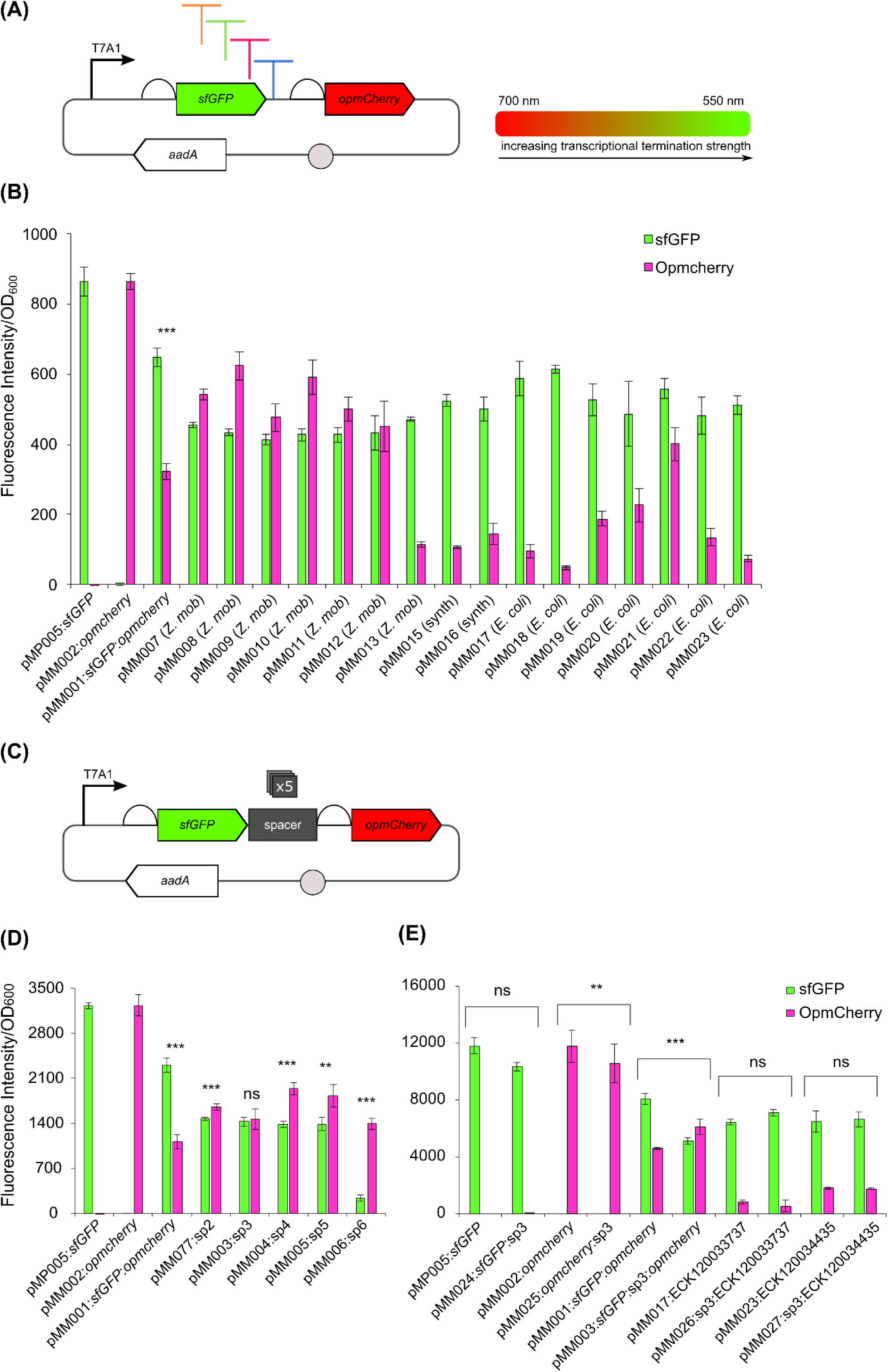
(A) Cloning of spacers in between *sfGFP* and *OpmCherry*. (B) Screening of spacers in *Z. mobilis*. The plasmids pMM077:spacer2, pMM003:spacer3, pMM004:spacer4, pMM005:spacer5 and pMM006:spacer6 were tested for an equal transcription of the *sfGFP* and *OpmCherry* reporters. The values reported were normalised by OD600. The following plasmids were used as controls: pMP005:*sfGFP*, pMM002:*OpmCherry* and pMM001:*sfGFP*:*OpmCherry*, which had previously demonstrated a strongly unbalanced transcription of the two reporters. Recombinant *Z. mobilis* cultures were grown in ZRMG rich medium at 30 °C, in a 96-well plate shaking at 800 rpm. *** = *p* < 0.01, ** = *p* < 0.05, ns = not significant, two-sample *t*-test. Error bars represent the standard deviation of three independent biological replicates. **(C) Final screening of terminators with and without spacer 3 in *Z. mobilis*.** Screening of the two strongest terminators, ECK120033737 and ECK120034435, and controls with and without spacer 3 using the *sfGFP* and *OpmCherry* dual-reporter system in *Z. mobilis*. The values reported are normalised by OD600. The recombinant *Z. mobilis* cultures were grown aerobically in ZRMG rich medium at 30 °C, in a 96-well plate, shaking at 800 rpm. Samples were harvested during late exponential phase and fluorescence measured using a microplate reader. *** = *p* < 0.01, ** = *p* < 0.05, ns = not significant, two-sample *t*-test. Error bars represent the standard deviation of three independent biological replicates.

To measure transcription termination efficiency (TE %), the sfGFP/OpmCherry fluorescence ratio of each dual-reporter system must be compared to a reference construct with equal reporters production (sfGFP/OpmCherry = 1), using the following formula:

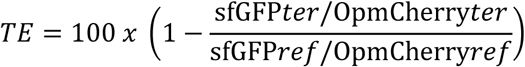

It was noticed however, that OpmCherry fluorescence was markedly lower than GFP fluorescence in strains containing the reference plasmid pMM001:*sfGFP*:*OpmCherry*. To obtain equal levels of fluorescence intensity, five short spacer DNA sequences (∼30-40 bp) lacking any complex secondary structures ^41^ were cloned between *sfGFP* and *OpmCherry* genes (Figure 5C and Table S4 in the supplemental information). The five spacer constructs were introduced into *Z. mobilis* and screened alongside control plasmids (Figure 5D). Among the tested sequences, spacer three yielded the most similar fluorescence levels between sfGFP and OpmCherry (two-sample *t*-test, *p* = 0.73), with a sfGFP/OpmCherry ratio of 1.05 and was used as the reference plasmid for subsequent terminator characterisation. This spacer sequence was also introduced onto the *sfGFP-* and *OpmCherry-*only control plasmids, and downstream of two of the strongest terminators ECK120033737 and ECK120034435 from the preliminary screening. The resulting plasmids introduced into *Z. mobilis* and rescreened for termination efficiency (Figure 5E).

No significant difference was observed between cultures containing the strong terminator plasmids lacking spacer 3 and those containing the plasmids with spacer 3 inserted (two-sample *t*-test between pMM017 and pMM026, *p* = 1; two-sample *t*-test between pMM017 and pMM026, *p* = 0.82). As the addition of a spacer did not alter the termination strength of ECK120033737 and ECK120034435, pMM003:*sfGFP*:sp3:*OpmCherry* was used as reference for measuring the TE of the previously screened terminator collection. The correlation of TE (%) and the Δ*G* of the predicted mRNA structures was tested, revealing a moderate inverse relationship (Pearson correlation coefficient *r* = -0.62) (Figure S7 in the supplemental information). Additionally, our data revealed a strong correlation between TE and increased sfGFP production (Figure S8 in the supplemental information; Pearson correlation coefficient *r* = 0.77), consistent with reports that terminators influence the stability of upstream mRNA sequences through secondary structure formation and protection from nuclease degradation ^42^.

### Bicistronic Designs for context-independent translation control

To date, tools to allow the predictable control of translation rates have not been developed for use in *Z. mobilis*. To tackle the limitations of RBSs context-dependency, bicistronic designs (BCDs) composed of two translationally coupled RBSs has been proposed. The first RBS mediates the strong translation of a short leader peptide (first cistron). Embedded in the leader peptide coding sequence is a second RBS that regulates the translation of a gene of interest (GOI) (second cistron). The sequence of the leader peptide overlaps by 1 base pair with the GOI, forming both a stop and a start codon via a -1 frame shift (Figure 6A). As the ribosome has an intrinsic helicase activity, it unwinds secondary structures that may have formed around the second RBS making it more accessible for translation, increasing the predictability of translation rates with different genetic constructs ^43^.

**Figure 6.**
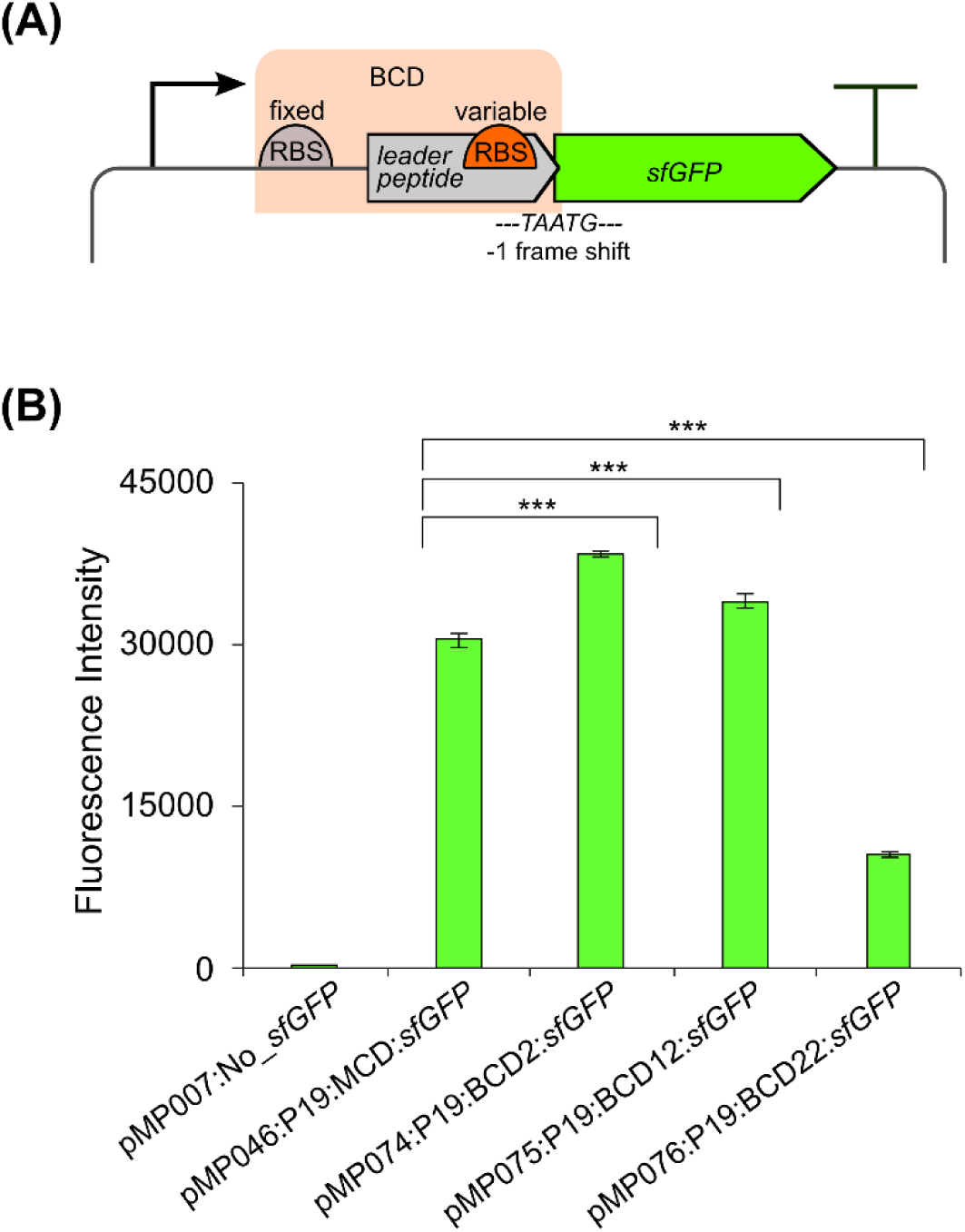
(A) Bicistronic design (BCD). The BCD is made of two-translationally coupled RBSs: a fixed RBS for strong translation of a short leader peptide (first cistron), and a variable RBS that controls the translation of a gene of interest (GOI) (second cistron). The stop codon of the leader peptide gene overlaps by 1 bp with the start codon of the GOI, causing a -1 frame shift that separates the translation of the two cistrons. The intrinsic helicase activity of the ribosome on the first cistron unwinds nearby secondary structures that may hinder ribosome binding to the second RBS. Therefore, the translation of the GOI becomes independent from the neighbouring nucleotide context. **(B) Screening of BCDs in *Z. mobilis*.** Three BCDs of different strengths -BCD2, BCD12 and BCD22- were screened in *Z. mobilis* by measuring the fluorescence of the downstream *sfGFP* reporter under the transcriptional control of the medium-strength promoter P19 (pMP074:19:BCD2:*sfGFP*, pMP075:P19:BCD12:*sfGFP*, pMP076:P19:BCD22:*sfGFP*). The empty plasmid pMP007:No_*sfGFP* and the plasmid pMP074:P19:*sfGFP* with a monocistronic RBS (100 k au), were used as controls. The recombinant *Z. mobilis* cultures were grown aerobically in ZRMG rich medium at 30 °C, in a 96-well plate shaking at 800 rpm. Samples were harvested during late exponential phase and fluorescence measured using flow-cytometry. *** = *p* < 0.01, a two-sample *t*-test was performed comparing the control pMP074:19:*sfGFP* with pMP074:19:BCD2:*sfGFP*, pMP075:P19:BCD12:*sfGFP* and pMP076:P19:BCD22:*sfGFP*. Error bars represent the standard deviation of three independent biological replicates.

From a set of twenty-two BCDs previously characterised in *E. coli* ^43^, a weak- (BCD22), medium- (BCD12) and high-strength (BCD2) BCD were selected for screening in *Z. mobilis* (Table S5 in the supplemental information). These were inserted between the medium-strength SPL promoter P19, and a gene encoding sfGFP. *Z. mobilis* transconjugants containing these three BCD-*sfGFP* constructs were screened by flow cytometry (Figure 6B). The three BCDs showed a strong, medium and weak pattern of fluorescence, in line with the observations reported in *E. coli*.

### Application of the Z. mobilis toolbox to the heterologous expression of a metabolic pathway for 2,3-butanediol

Having established a broad set of constitutive promoters with a wider expression range than previously reported, along with strong transcriptional terminators, we could now use this toolkit to rapidly assemble and explore the design space of a multi-enzyme metabolic pathway in *Z. mobilis* for the first time. We chose the biosynthesis of 2,3-butanediol (2,3-BDO), a platform chemical with broad applications in fuels, solvents, synthetic rubbers, pharmaceuticals, and cosmetic products ^44^ for this proof-of-concept. The pathway consists of three enzymes: acetolactate synthase (encoded by *als*); acetolactate decarboxylase (encoded by *aldC*); and butanediol dehydrogenase (encoded by *bdh*) ^20^ (Figure 7A).

**Figure 7.**
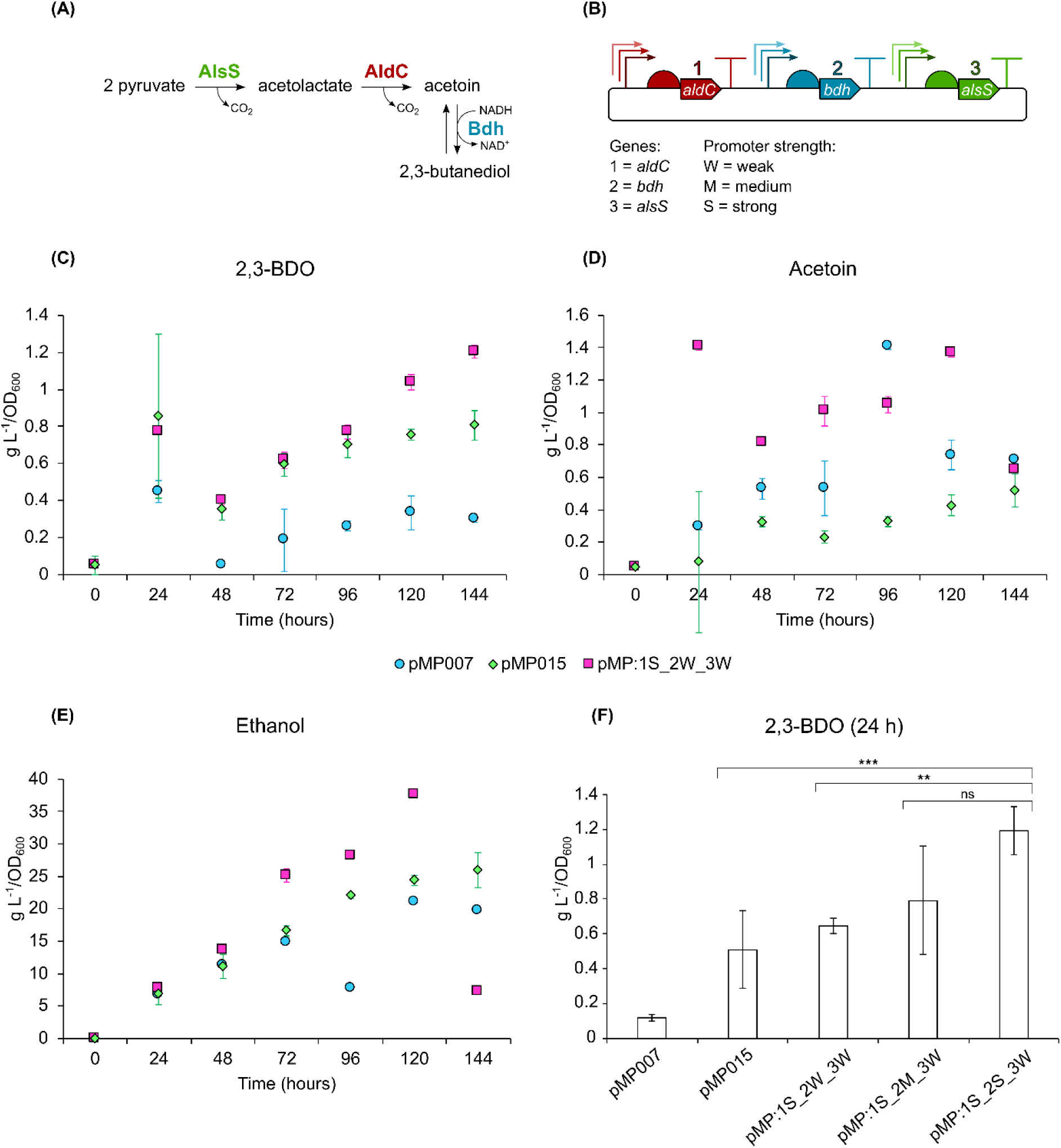
(A) Pathway for the biosynthesis of 2,3-butanediol (2,3-BDO). Two molecules of pyruvate are combined by the acetolactate synthase (AlsS) into acetolactate with the release of one CO2. Acetolactate is decarboxylated by the acetolactate decarboxylase (AldC) into acetoin, which is reduced by the butanediol dehydrogenase (Bdh) into 2,3-butanediol, with the formation of NAD+. The last reaction from acetoin to 2,3-butanediol is reversible. **(B) Combinations of three different weak, medium and strong promoters upstream of *aldC*, *bdh* and *alsS*, yield up to twenty-seven differentially regulated 2,3-BDO pathways.** When referring to the plasmids carrying the different 2,3-BDO pathways, we used pMP followed by the number associated to each gene (1 = *aldC*, 2 = *bdh*, 3 = *alsS*) and a letter indicating the strength of the promoter upstream of that gene (W = weak, M = medium, S = strong). **(C), (D) and (E) respectively show the 2,3-BDO, acetoin and ethanol production of *Z. mobilis* carrying the negative control pMP007 (blue circles), the positive control pMP015 (green diamonds) and pMP:1S_2W_3W (pink squares).** Cultures were grown in ZRMG80 minimal medium for six days, and samples for HPLC analysis harvested every 24 hours. The metabolites concentration was normalised by OD600. Error bars represent the standard deviation of three independent biological replicates. **(F) 2,3-BDO production of the negative and positive controls *Z. mobilis* (pMP007), (pMP015) respectively, and *Z. mobilis* (pMP:1S_2W_3W), (pMP:1S_2M_3W) and (pMP:1S_2S_3W).** Cultures were grown in ZRMG rich medium, and samples harvested after 24 hours of growth. 2,3-BDO was measured by NMR, and concentrations were normalised by OD600. Error bars represent the standard deviation of three independent biological replicates. *** = *p* < 0.01, ** = *p* < 0.05, ns = not significant, two-sample *t*-test.

The 2,3-butanediol (2,3-BDO) pathway has been previously introduced into *Z. mobilis* 9C by expressing the *aldC* and *bdh* genes from *Enterobacter cloacae*, and *alsS* from *Bacillus subtilis*^20^. The three genes and their cognate RBSs were carried on the plasmid pEZ-BC11, with expression of *aldC* and *bdh* driven by the native P*gap* promoter and *alsS* controlled by the inducible P*tet* promoter, however left intentionally uninduced ^20^. To allow independent control of transcription for each gene in the pathway, and for the rapid combinatorial assembly of regulatory elements and genes, the Start-Stop one-pot assembly method was used (Figure S9 and S10 in the supplemental information) ^45^. The pathway-encoding genes, nine promoters (three weak, three medium and three strong promoters) and the three strongest terminators were cloned into a level 0 holding vector. Level 1 assemblies were carried out with each gene being combined with a weak, medium or strong promoter (Table 1). A level 2 expression vector, pMP081, was constructed for faithful propagation in *Z. mobilis* and a mix of all nine level 1 plasmids used for the final level 2 plasmid assembly, generating a library of twenty-seven combinations of differentially regulated 2,3-BDO pathway genes (Figure 7B). Finally, as 2,3- BDO had been successfully produced in *Z. mobilis* 9C using pEZ-BC11 ^20^, the same promoter-gene layout was reproduced in the construction of the control plasmid pMP015.

**Table 1.**
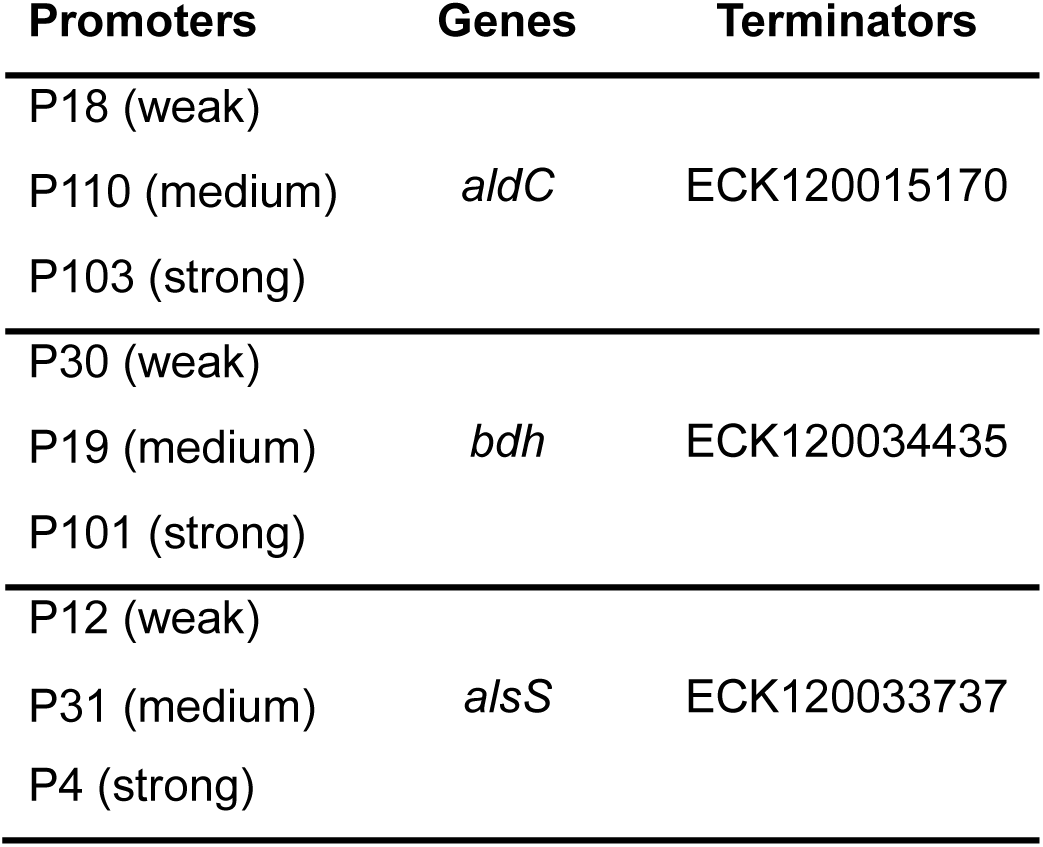
Promoters, genes and terminators used to build combinatorial 2,3-BDO pathways.

The level 2 plasmid library was introduced into *Z. mobilis*, and thirteen colonies arbitrarily selected for plasmid sequencing to assess assembly efficiency and fidelity. All thirteen plasmids had unique and complete 2,3-BDO pathways. Sequenced plasmids were renamed pMP, followed by numbers for each gene of the pathway (1 = *aldC*, 2 = *bdh*, 3 = *alsS*) coupled to letters indicating promoter strength (W = weak, M = medium, S = strong). To screen for 2,3- BDO production, the supernatants from thirty individual *Z. mobilis* transconjugant cultures, consisting of a mixture of sequenced and unsequenced plasmids, were assessed by HPLC coupled to refractive index detection, with *Z. mobilis* (pMP015) and (pMP007) as positive and negative controls respectively (Figure S11 in the supplemental information). As rich medium was unsuitable for our HPLC analysis due to the high background absorbance and refractive interference, ZMMG80 minimal medium was used to culture *Z. mobilis* transconjugants for six days. A wide range of 2,3-BDO yields were observed across the library, with four combinatorial assembly strains showing similar yields to the positive control strain and one showing higher 2,3-BDO levels. These five strains were selected for further characterisation (Table 2).

**Table 2.** List of plasmids carrying differentially regulated 2,3-BDO pathways that were selected for further characterisation.

|  | 1) *aldC* promoter | 2) *bdh* promoter | 3) *alsS* promoter |
| --- | --- | --- | --- |
| **pMP:1S_2W_3W** | P103 (strong) | P30 (weak) | P12 (weak) |
| **pMP:1M_2W_3S** | P110 (medium) | P30 (weak) | P4 (strong) |
| **pMP:1M_2S_3M** | P110 (medium) | P101 (strong) | P31 (medium) |
| **pMP:1S_2W_3S** | P18 (weak) | P19 (medium) | P12 (weak) |
| **pMP:1W_2M_3W** | P103 (strong) | P30 (weak) | P4 (strong) |

Cultures containing these plasmids were grown in triplicate for six days, with samples collected every 24 hours and supernatants analysed by HPLC for 2,3-BDO, acetoin and ethanol production (Figure 7C, 7D, 7E, and Figure S12 in the supplemental information). In line with the preliminary screening, the best 2,3-BDO-producing strain was *Z. mobilis* pMP:1S_2W_3W with 1.20 ± 0.034 g L^-1^/OD600 of final product, while the control (pMP015) yielded 0.80 ± 0.078 g L^-1^/OD600 of 2,3-BDO (Figure 7C). Only one strain was tested with a strong promoter located upstream of *bdh*, *Z. mobilis* pMP:1M_2S_3M, and as it accumulated acetoin levels that were significantly lower than those of the top 2,3-BDO producer, *Z. mobilis* pMP:1S_2W_3W, across five of the six days of culture (two sample *t*-test, Day 1 *p* = 0.022, Day 2 *p* = 0.035, Day 3 *p* = 0.004, Day 4 *p* = 0.01, Day 5 *p* = 0.004, Day 6 *p* = 0.068) (Figure S13 in the supplemental information), we hypothesised that a stronger promoter upstream of *bdh* on pMP:1S_2W_3W could increase the conversion of acetoin to 2,3-BDO. To test this, we designed pMP:1S_2M_3W and pMP:1S_2S_3W, carrying the medium-strength promoter P19 and strong promoter P101, respectively. To allow culturing and testing in rich media, the supernatant of larger cultures was tested using NMR instead of HPLC. The supernatants of strains containing pMP:1S_2M_3W or pMP:1S_2S_3W were tested after 24 hours of growth (Figure 7F) and the rationally designed *Z. mobilis* (pMP:1S_2S_3W) outperformed both the reference strain (pMP015) and the top 2,3-BDO producer from the initial screen (pMP:1S_2W_3W) (two-sample *t*-test, *p* = 10^-4^, *p* = 0.018, respectively). In line with our rationale, this increase in 2,3-BDO corresponded to a reduction in acetoin accumulation in the best 2,3-BDO producer (Figure S14 in the supplemental information).

## Discussion

In this work, we provide the most comprehensive genetic toolkit for the engineering of *Z. mobilis* to date. Although the final set of 39 promoters featured non-standard architectures, including operator sequences lacking highly conserved nucleotides and, in some cases, shorter sequences, these promoters still resulted in a wide range of expression strengths. Crucially, many strong promoters were identified with expression levels much higher than the previously reported “strongest” constitutive promoter, P*gap*. This wider range of expression strengths will allow multi-enzyme pathways such as metabolic pathways to be constructed, where the level of each enzyme can be modulated with unprecedented resolution, using high-throughput one-pot assembly methods.

Reintroducing the most conserved nucleotides to three of the synthetic promoters also resulted in transcription, suggesting that the sigma 70 (σ⁷⁰) factor of RNA polymerase in *Z. mobilis* can bind and initiate transcription at a much more diverse sequences than previously thought. Indeed, recent work suggests that *de novo* promoters emerge much more frequently in random DNA than genomic DNA ^46^, suggesting perhaps that design strategies for synthetic promoters could start from sequences that do not resemble natural promoters at all.

When the activity of the same promoters was compared between both *Z. mobilis* and *E. coli*, there was poor agreement between the two organisms relative to a 1:1 line. Indeed, the promoters chosen through screening of the random library to obtain a nice range of strengths in *Z. mobilis*, showed a limited range of strengths in *E. coli.* This highlights the importance of host-tailored promoter selection for expression. The diverse transcriptional strength may be attributed to different σ⁷⁰ domains in *Z. mobilis* and *E. coli*. Notably, when comparing the amino acid sequence of the four σ⁷⁰ domains in the two organisms, there are high degrees of similarity in the σ^2^ domain (100% pairwise identity), σ^3^ (74% pairwise identity) and σ^4^ (83% pairwise identity), whereas the σ^1^ domain has only 29% pairwise identity. It is possible that these differences are sufficient for a diverse interaction of the σ⁷⁰ with the promoters, leading to a significant change in strength between hosts. This lack of equality of expression can however be used as a strength, allowing the cloning of genes that produce products that may be toxic in *E. coli*, before being tested in *Z. mobilis*, lowering transcriptional and translational burden, and reducing selective pressure to mutate sequences during plasmid propagation.

As fermentation and the production of highly-reduced chemicals is one of the main benefits of *Z. mobilis* over other bacterial hosts, we investigated for the first time promoter performance under anaerobic conditions. Since metabolism varies with oxygen availability, different proteins, including transcription factors, may be produced anaerobically, potentially influencing transcription initiation. Our results suggest that the synthetic promoter library is not affected by the absence of oxygen, something not commonly tested in synthetic biology toolkit studies.

To achieve controlled insulation of gene expression in *Z. mobilis*, seven predicted native intrinsic terminators were tested. Surprisingly, six of the seven native terminators showed no transcriptional termination, suggesting that *Z. mobilis* terminators may differ from the standard stem-loop and poly-U motifs on which prediction tool (FindTerm^39^) was trained. In line with this, a recent multispecies comparisons of intrinsic terminators revealed substantial differences in nucleotide composition, highlighting the limitations of computational tools for cross-species prediction ^47^. Importantly, strong heterologous and synthetic terminators were identified, providing effective tools for pathway orthogonality and rational engineering of complex metabolic pathways in *Z. mobilis*.

The synthetic promoters and strong terminators developed and characterised here were made compatible with the high-throughput Start–Stop assembly method, allowing modular and scarless cloning of both coding and non-coding parts. This approach was used to rapidly construct and screen various promoter and 2,3-BDO pathway gene combinations, which were then further refined through rational engineering to boost production. This demonstrated how integration of combinatorial design with targeted optimisation can accelerate the development of high-performing strains. No metabolic engineering was carried out to optimise 2,3-BDO yields further, but this would be a promising next step for commercialisation.

## Concluding remarks

Engineering in *Z. mobilis* has been mostly limited to the use of poorly characterised regulatory parts. Here, we developed a novel molecular toolbox including an extensive synthetic promoter library (SPL), context-independent bicistronic translational regulators (BCDs), and robust intrinsic terminators. By integrating combinatorial and rational design, this toolbox was able to modulate the expression of the heterologous 2,3-BDO pathway, generating a top 2,3- BDO producer *Z. mobilis* strain. From this proof of concept, we demonstrated that these regulatory elements open new opportunities for predictable pathway optimisation and broader use of *Z. mobilis* as host for industrial biotechnology purposes. Our toolkit vastly expands the suite of molecular tools available for predictable and precise molecular engineering in this organism.

## Materials and Methods

### Bacterial strains and culture conditions

The high-transformation efficiency *Z. mobilis* ZM4 (*hsdSc hsdSp*) strain, kindly gifted by Professor Patricia Kiley from University of Wisconsin Madison, *E. coli* TOP10, and the diaminopimelic acid auxotrophic *E. coli* WM6026 were used in this study. For simplicity, *Z. mobilis* ZM4 (Δ*hsdSc* Δ*hsdSp*) is referred to as *Z. mobilis* throughout the text.

*Z. mobilis* was grown in Rich Medium Glucose (ZRMG), containing 20 g/L glucose, 2 g/L KH2PO4, 10 g/L yeast extract (pH 6-6.5), and Minimal Medium Glucose (ZMMG), containing a base solution of 1 g/L KH2PO4, 1 g/L K2HPO4, 500 mg/L NaCl, 1 g/L (NH)4SO4, and a trace elements solution of 200 mg/L MgSO4 x 7H2O, 0.010 g/L CaCl2 x 2H2O, 25 mg/L Na2MoO4 x 2H2O, 25 mg/L FeSO4 x 7H2O, 1 mg/L Calcium pantothenate (pH 6-6.5). Liquid cultures of *Z. mobilis* strains were grown at 30°C statistically or shaking at 120 rpm. *E. coli* strains were grown in rich Luria Bertani (LB) medium. When culturing *E. coli* WM6026, a 0.1 mM final concentration of diaminopimelic acid (DAP) was added to the medium. Liquid cultures of *E. coli* strains were grown at 37°C shaking at 200 rpm.

### Conjugation

Conjugation was performed to transfer plasmids from *E. coli* WM6026 (donor strain) to *Z. mobilis* (recipient strain). *E. coli* WM6026 was grown in 5 ml LB medium supplemented with 0.1 mM DAP shaking at 37°C, and *Z. mobilis* was grown in 5 ml ZRMG static at 30°C. At a similar OD600 of ∼0.5, 1 ml of each culture were combined and spun down for 30 seconds at 15,000 x *g*. The supernatant was partially discarded, and the pellet resuspended. The cell suspension was placed as a single drop on a ZRMG plate supplemented with 10 g/L tryptone and 0.1 mM DAP, to allow the mating of the donor and recipient overnight. The next day, the cells were resuspended in ZRMG and grown for 2 hours static at 30°C, then plated on ZRMG supplemented with the appropriate antibiotic.

### Construction of plasmids for the synthetic promoter library

The plasmids used in this chapter are derived from pPK15404, which was kindly gifted by Professor Patricia Kiley from University of Wisconsin Madison ^53^. pPK15404 contains the *E. coli lacZ* gene, pBBR-1 origin of replication, the *mob* gene and spectinomycin resistance (*aadA*). Using Gibson assembly, the *lacZ* gene was replaced by an expression unit including the terminator ECK120010799 ^54^, T7A1 and *sfGFP*, generating pMP005. This was used as template for building all the synthetic promoter library constructs, where the T7A1 promoter was replaced by inverse PCR with P*gap*_long (pMP003), P*gap*_short (pMP004), and partially randomised promoters. These were cloned using degenerate primers with tails carrying the following sequence: 5’ - (N)10TAWAAT(N)17TTGMHD(N)6 - 3’, in which “N” could be any nucleotide, “W” an adenine or thymine, “M” an adenine or cytosine, “H” an adenine, a cytosine or a thymine, and “D” an adenine, a guanine, or a thymine.

### Construction of plasmids for anaerobic testing of synthetic promoters

The *Y-FAST* gene was isolated from the plasmid pCK871 and cloned by Gibson Assembly downstream of the T7A1 in pMP005, the P*gap*_long in pMP003, the P*gap*_short in pMP004. The low, medium and high-strength promoters P12, P19 and P103 were introduced upstream of the *Y-FAST* using inverse PCR on the template pMP068:T7A1:*Y-FAST*.

### Construction of plasmids for the terminator library

pMP005 was used as template for building the dual-reporter system with the T7A1 promoter, *sfGFP* and *OpmCherry*. *OpmCherry* and its RBS were isolated from pEZ15A ^26^ and introduced downstream of the *sfGFP* in pMP005 by Gibson Assembly, creating pMM001. This was the template for cloning terminators and spacers, which were introduced by inverse PCR between the *sfGFP* and *OpmCherry*.

### Construction of plasmids for the BCDs characterisation

The plasmid pMP046, carrying the medium-strength synthetic promoter P19 upstream of the *sfGFP* was used as the basis for the cloning and testing of BCD2, BCD12 and BCD22. Using Gibson Assembly, P19 and BCD22 were cloned in pMP005, in place of T7A1 and the monocistronic RBS upstream the *sfGFP*, generating pMP076. BCD2 and BCD12 were introduced by inverse PCR in pMP076 in place of BCD22.

### Preparation of cells for fluorescence measurements

In a 96-deep well plate, biological triplicates of single colonies were inoculated in a 300 µl of rich medium and grown overnight. The next day, the cultures were diluted 1:100 in 150 µl of rich medium in a 96-well plate and grown until mid-exponential phase (0.4-0.6 OD600). At this stage the cultures were subsequently diluted to an OD600 of 0.05 in 150 µl of rich medium in a 96-well plate and grown to mid-exponential phase (0.8 – 1.0 OD600). The plate reader and the flow-cytometer were used to read the fluorescence of the cells in this phase of growth. Prior to the use of the flow-cytometer, the cells were diluted 1:200 in filtered phosphate buffered saline (PBS) in a 96 well-plate. When measuring the fluorescence of the Y-FAST reporter an 20 µM quantity of chromogen was added to the resuspension of cells in PBS.

### Measurement of fluorescence using the plate reader

Using the Plate Reader Spark (Tecan), *sfGFP* was measured at 485 nm excitation wavelength and 535 nm emission wavelength, OpmCherry was measured at 560 nm excitation wavelength and 620 nm emission wavelength. The fluorescence of the cells was measured in a 96-well plate, shaking at 800 rpm or static depending on aerobiosis or anaerobiosis growth, with an interval of readings of 15 minutes. The temperature was set at 30°C and 37°C when respectively growing *Z. mobilis* and *E. coli*. The fluorescence data reported in this chapter were collected during the late exponential growth phase.

### Flow-cytometry

The flow-cytometer FACSCanto™ II was used for measuring the fluorescence of the synthetic promoter library. For *sfGFP* detection, a laser with a 450 V was used, while size and granularity of the cells were measured with the Forward Scatter (FSC) and Side Scatter (SSC) lasers, respectively using 413 V and 334 V. The sample volume injected was 50 µl, and the sample flow rate was 1.5 µl/sec. The machine was set to record 10,000 events for each sample. Data were analysed using FCS express software (De Novo Software), and the geometric means of the population obtained.

### Start-Stop Assembly reaction

Start-Stop Assembly reactions contained 20 fmol of destination vector plasmid DNA, 40 fmol of each insert, 2 µl T4 buffer, 1 µl T4 ligase, 1 µl SapI/BsaI in a 20 µl reaction. Reactions were incubated for 30 cycles at 37 °C for 5 minutes and at 16 °C for 5 minutes, followed by a final stage at 65 °C for 20 minutes.

### Construction of 2,3-butanediol plasmids

Up to twenty-seven combinations of 2,3-BDO pathways were created following the Start-Stop system for DNA Assembly as described by Heap *et al*. ^45^

### Level 0

The vector pGT400 was used as level 0 to store individual parts, including promoters, open reading frames and terminators. Back-to-back primers with overhangs were used to introduce promoters and terminators in pGT400, while open-reading frames were introduced into pGT400 using Gibson Assembly. The level 0 cargoes were flanked by two inward-facing SapI restriction sites, which generated specific fusion sites for their assembly into level 1 vectors.

### Level 1

The Start-Stop assembly reaction was conducted on the level 0 vectors and the level 1 vectors creating full expression units through SapI digestion and T4 ligation, as previously described^45^. pGT402, pGT404 and pGT405 were used as level 1 destination vectors. The level 1 vectors are reported in . These are characterised by two outward-facing SapI sites and two inward-facing BsaI sites. The BsaI sites generate cohesive ends that enable a sequential cloning of the expression units in whole pathways in a level 2 vector.

### Level 2

A lacZ gene flanked by two outward-facing BsaI sites was isolated from pGT417 and cloned via Gibson Assembly in the pMP002 backbone creating the level 2 plasmid pMP081. pMP081 carried the A and Z fusion sites to allow the ligation of the whole plasmids into its backbone.

### HPLC samples preparation and measurement

Every 24 h, 1 ml of samples were taken from the *Z. mobilis* cultures for OD600 measurement and to evaluate the concentration of 2,3-BDO, ethanol and acetoin using HPLC (High-Performance Liquid Chromatography). Samples were stored at -4 C° until the end of bacterial growth (144h), when the samples were prepared for injection in the HPLC. To remove bacterial cells, samples were transferred through a 0.2-μm filter into HPLC vials. Concentrations of 2,3- BDO, acetoin and ethanol were determined by Agilent 1100 series HPLC (Agilent, CA) utilizing a Bio-Rad Aminex HPX-87H organic acids column and Cation H^+^ guard cartridge (Bio-Rad, CA) operating at 65 °C. A refractive index detector was used for compound detection. 0.05 mM sulfuric acid was used as the isocratic mobile phase at a flow rate of 0.6 mL/min ^55^.

### NMR

NMR Spectroscopy was performed using a JEOL ELS600 (6.0.0) Delta Spectrometer at frequencies of 400 MHz for *1*H NMR and 101 MHz for *13*C NMR. Chemical shifts (δ) were recorded in parts per million (ppm) and were relative to an internal standard, such as tetramethyl silane (TMS). Deuterated solvents used included deuterated dimethyl sulfoxide (DMSO-*d6*), deuterated chloroform (CDCl*3*), deuterated water (D*2*O) and deuterated methanol (CD*3*OD).

## Acknowledgments

This work was funded by the EPSRC ReNU (Renewable Energy Northeast Universities) CDT at Northumbria University.

