## Supplemental Information for "A Comprehensive Genetic Toolkit for Rapid and High-Resolution Engineering of Metabolic Pathways in *Zymomonas mobilis*"

### Supplementary Information for: A comprehensive genetic toolkit for *Zymomonas mobilis*; allowing rapid, high-resolution, combinatorial engineering of metabolic pathways.

**(A)** -35 box

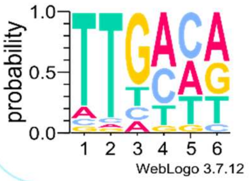


| | 1 | 2 | 3 | 4 | 5 | 6 |
|---|---|---|---|---|---|---|
| A | 11.04% | 4.56% | 10.94% | 54.27% | 31.71% | 41.76% |
| T | 77.86% | 88.17% | 17.16% | 15.12% | 26.16% | 25.80% |
| C | 5.97% | 4.03% | 9.79% | 27.05% | 35.27% | 7.53% |
| G | 5.13% | 3.24% | 62.11% | 3.56% | 6.86% | 24.91% |

**(B)** -10 box

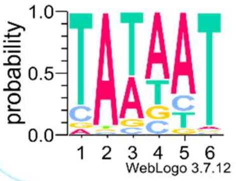


| | 1 | 2 | 3 | 4 | 5 | 6 |
|---|---|---|---|---|---|---|
| A | 4.60% | 92.05% | 37.10% | 59.60% | 67.77% | 2.83% |
| T | 78.81% | 2.93% | 50.90% | 19.15% | 14.39% | 93.77% |
| C | 11.72% | 1.78% | 4.70% | 9.79% | 13.24% | 1.62% |
| G | 4.87% | 3.24% | 7.30% | 11.46% | 4.60% | 1.78% |

**Figure S1. Nucleotide composition of the −35 and −10 boxes of *Z. mobilis* putative $\sigma^{70}$ promoters. (A)** The left panel shows the nucleotide frequency at each position of the −35 box (A = adenine, T = thymine, C = cytosine, G = guanine), visualized as a sequence logo generated using WebLogo. The right panel displays the percentage composition of each nucleotide per position. Highlighted cells indicate the nucleotide(s) that have a ≥ 60% frequency, or, when this threshold is not met, the nucleotide(s) with the highest percentage, and those with the next greatest value with less than 5% difference between them. **(B)** The same analysis and visualisation approach were applied to the −10 box.

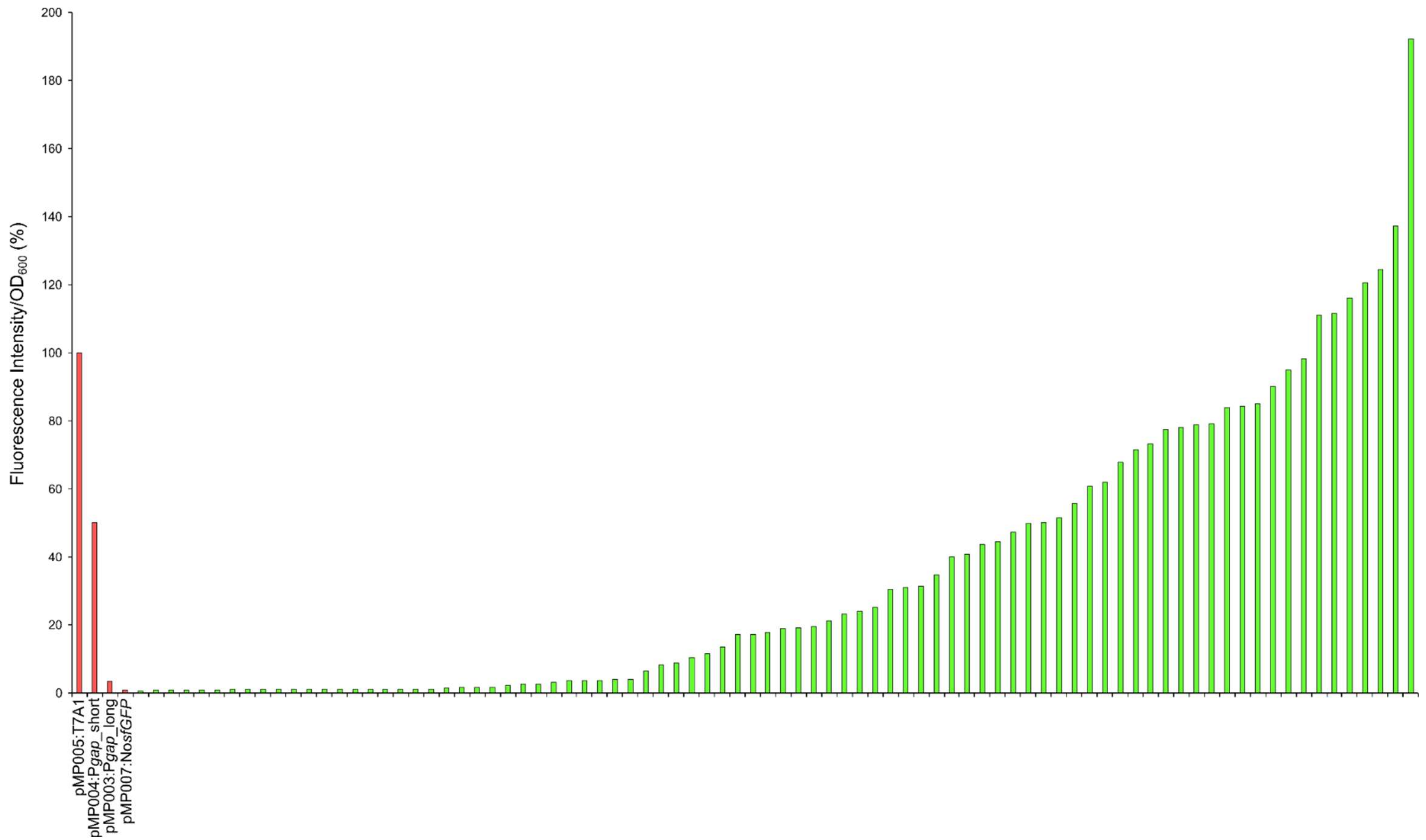


**Figure S2. Preliminary screening of fluorescence distribution of synthetic promoters in *Z. mobilis*.** Eighty-four colonies arbitrarily chosen from a plate were tested along the following constructs: pMP005:T7A1, pMP003:P*gap*_long, pMP004:P*gap*_short and pMP007:No_*sfGFP*. The fluorescence is shown in percentage relative to the positive control T7A1 (100%). Fluorescence was measured using the plate reader. *Z. mobilis* recombinant strains were grown aerobically in ZRMG rich medium, at 30°C, in 96-well plates, shaking at 800 rpm. Samples were harvested during late exponential phase.

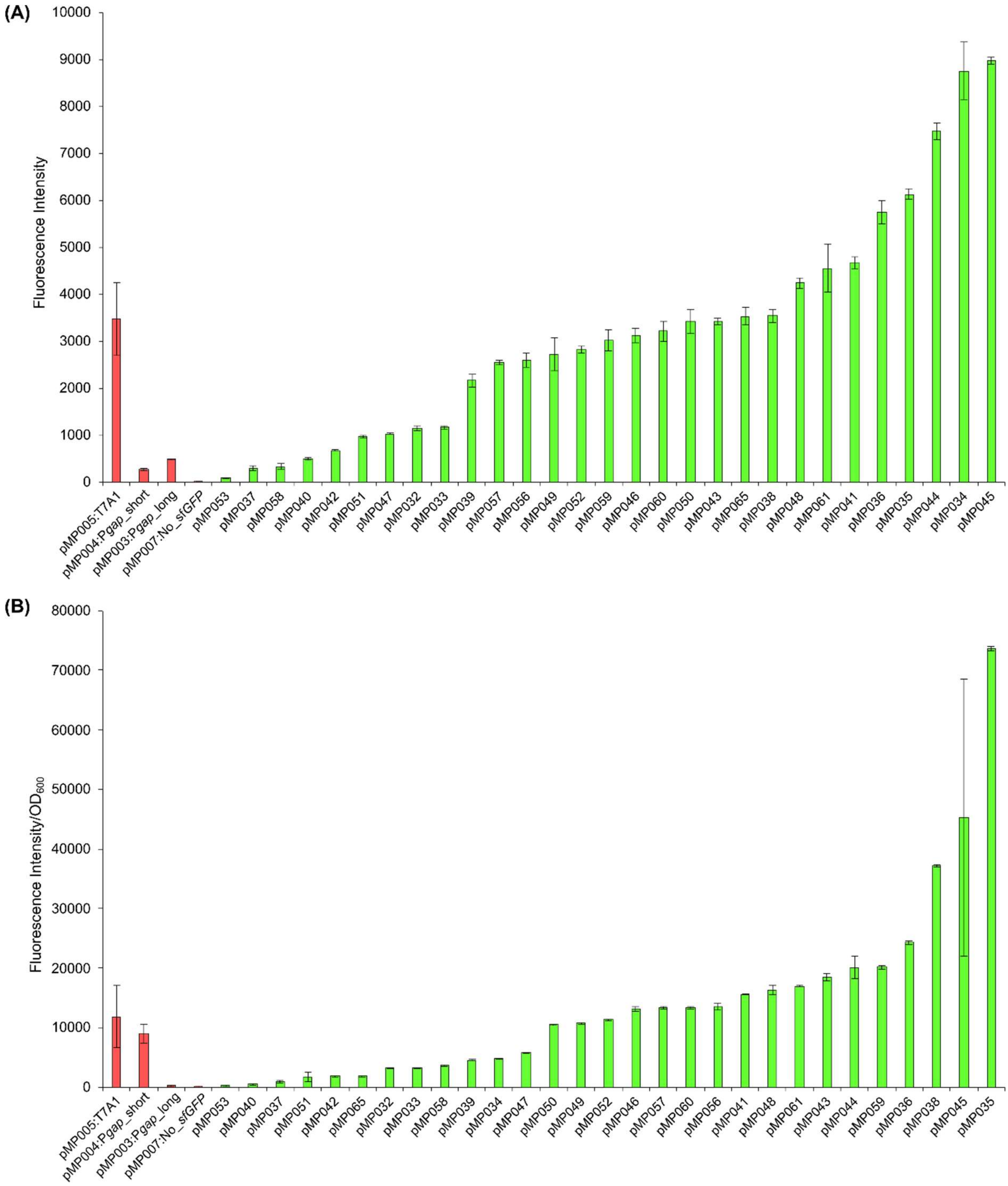


**Figure S3. Screening of fluorescence distribution of synthetic promoters in *Z. mobilis* (A) and *E. coli* WM6026 (B)**. Twenty-nine synthetic promoters from the preliminary screening having at least 2% difference in *Z. mobilis* were further screened in triplicate by flow-cytometry. The selected recombinant colonies were tested along with the following controls: pMP005:T7A1, pMP003:P*gap*_long, pMP004:P*gap*_short and pMP007:No_*sfGFP*. *E. coli* WM6026 and *Z. mobilis* recombinant strains were grown aerobically in rich medium, respectively in LB at 37°C 70 and in ZRMG at 30°C, in 96-well plates, shaking at 800 rpm. Samples were harvested during late exponential phase. Error bars represent the standard deviation of three independent biological replicates.

**Table S1. Expression groups following post-hoc Welch pairwise tests with Benjamini–Hochberg correction.**

| Promoter | Mean Fluorescence (geometric mean, a.u.) | CLD |
|---|---|---|
| pMP045 | 8977.275951 | G |
| pMP034 | 8775.256392 | E |
| pMP044 | 7470.871301 | FTUVW |
| pMP035 | 6127.393537 | FG |
| pMP036 | 5750.64995 | H |
| pMP041 | 4674.034902 | R |
| pMP061 | 4566.489282 | IMPS |
| pMP048 | 4240.599139 | A |
| T7A1 | 3538.745462 | HIMOPQSUWXZEGJLN |
| pMP038 | 3534.813405 | JK |
| pMP065 | 3528.637433 | OPWG |
| pMP043 | 3421.092291 | L |
| pMP050 | 3416.519318 | NOVWYZBCDE |
| pMP060 | 3214.509778 | OWZGJ |
| pMP046 | 3117.272375 | T |
| pMP059 | 3032.979669 | NVYDFIKM |
| pMP052 | 2808.379646 | B |
| pMP049 | 2728.361823 | MQUX |
| pMP056 | 2592.901588 | CH |
| pMP057 | 2548.830544 | DE |
| pMP039 | 2159.11881 | LMNOP |
| pMP033 | 1157.8302 | D |
| pMP032 | 1134.433067 | C |
| pMP047 | 1029.670052 | DXYZ |
| pMP051 | 971.5054456 | FG |
| pMP042 | 679.4177376 | S |
| pMP040 | 499.7997358 | AQ |
| Long_Gap | 493.6661934 | NA |
| pMP058 | 327.256191 | BJRKL |
| pMP037 | 288.9603739 | I |
| Short_Gap | 264.0551157 | KMN |
| pMP053 | 91.73297141 | HIJ |
| No_EGFP | 18.55078339 | AB |

**Table S2. Synthetic promoter library.** The table reports the name of the twenty-nine synthetic promoters with their corresponding plasmid, their nucleotide sequence and their length. The promoters are ranked based on increasing transcriptional strength.

| Plasmid | Promoter | Promoter sequence (5'-3') | Length |
|---|---|---|---|
| pMP032 | P29 | CCGACCAACTTAAAATGATGGAGTTCATGTTGCTTGTTTAG | 41 nt |
| pMP033 | P78 | CCGACCAACTTAAAATGATGGAGTTCATGTTGCTTGCTTAGCAAG | 45 nt |
| pMP034 | P101 | CCCCGTATGCTATAATATGCTGGAGAGCTGTTCTTGATGCATGTG | 45 nt |
| pMP035 | P111 | CGCATCTGAATATAATTGGGGTGACTTTTATTGAAAACATTC | 42 nt |
| pMP036 | P109 | AACATGCTCTTATAATGATTGGGTTTAGTTTGGTTGACGTAATTC | 45 nt |
| pMP037 | P18 | CTTGACCACATAAAATTAAGCAATGCTGGGCGATTGACGTAGATT | 45 nt |
| pMP038 | P112 | TCCCCGCAAATATAATGATAGCTAGCGAAGGTGTTGACGTAGCAC | 45 nt |
| pMP065 | P17 | TAGGGGAAGATAAAATACATCACAGCTTTAATCTTGACGACTGGT | 45 nt |
| pMP039 | P113 | AGAACACTAATAAAATAGATAAGTGGTGTAGGCTTGATTTCTAGG | 45 nt |
| pMP040 | P12 | TCATGATTAATATAATTAC | 18 nt |
| pMP041 | P115 | CTAGATACAATAAAATTGTGAAGCCTACAACCGTTGATATCCTGG | 45 nt |
| pMP042 | P25 | TTAGAGGAAATAAAATTGAATCGCACGATCTATTTGCATTATGCT | 45 nt |
| pMP043 | P11 | CACAGTACTATAAAATTAACCTGTTGATTGTAGAA | 35 nt |
| pMP044 | P4 | GTAAATACTATATAATAACTGATTATTGCTTATTGTT | 37 nt |
| pMP045 | P103 | ACAACAATAGTATAATGTTCGAACTATAATCCTTGCAGTGTTTT | 44 nt |
| pMP046 | P19 | CGGAAGACACTAAAATAAAGACCTTTGAATTTTACG | 36 nt |
| pMP047 | P116 | GTCTATAACTTAAAATGTGAGGCCTATAAACTTTTGCTAATCGCT | 45 nt |
| pMP048 | P6 | TGATCAACATTAAAATAGGGCAAGTTCTCGAAATTGCAGTCAGTA | 45 nt |
| pMP049 | P28 | TCGTAAGCATTAAAATCCTGAAATTGACTTCAAAT | 35 nt |
| pMP050 | P82 | GGTATAAGCGTAAAATATCAATACTAAGGGATATTGAAGTCTCAT | 45 nt |
| pMP051 | P67 | CCTACGCGCGTAAAATCGGCAAAGTAGGTTTTATTGCTGCCCTAT | 45 nt |
| pMP052 | P31 | AAACCGAGAATATAATTATACAGATATTTCCGATTGCTGCCCTAT | 45 nt |
| pMP053 | P15 | AGAGCATTCATATAATGATCACTATAGTTTATTGCCATTCCAG | 43 nt |
| pMP054 | P16 | AGCCTGTAATTAAAATACTGTTCTTGACGAGTTGC | 35 nt |
| pMP055 | P15 | AAGATGTACTTAAAATAACGAAATCTTCTGATTTTGCCTGAGTAG | 45 nt |
| pMP056 | P8 | CTTAAATAACTATAATTCGAACATCAATGTGTGTTGCAAACGACA | 45 nt |
| pMP057 | P108 | CATATCACTGTATAATAATGAGATTGAGGTGTGTTGCCTTATCGG | 45 nt |
| pMP058 | P30 | GACGACAATTGGCTGGGAACGGTATACTGGGAATAAATGG | 39 nt |
| pMP059 | P110 | CGAAAGCCGATAAAATAATCCCGGAATAAATGG | 33 nt |
| pMP060 | P107 | TTCCCTGACGTATAATCCCGTGATCAGTTTTTGTTGACGAGAA | 43 nt |
| pMP061 | P106 | CGATAAGCGGTATAATGCCATATCGGGCAATTGTTGCCGAGGTGG | 45 nt |

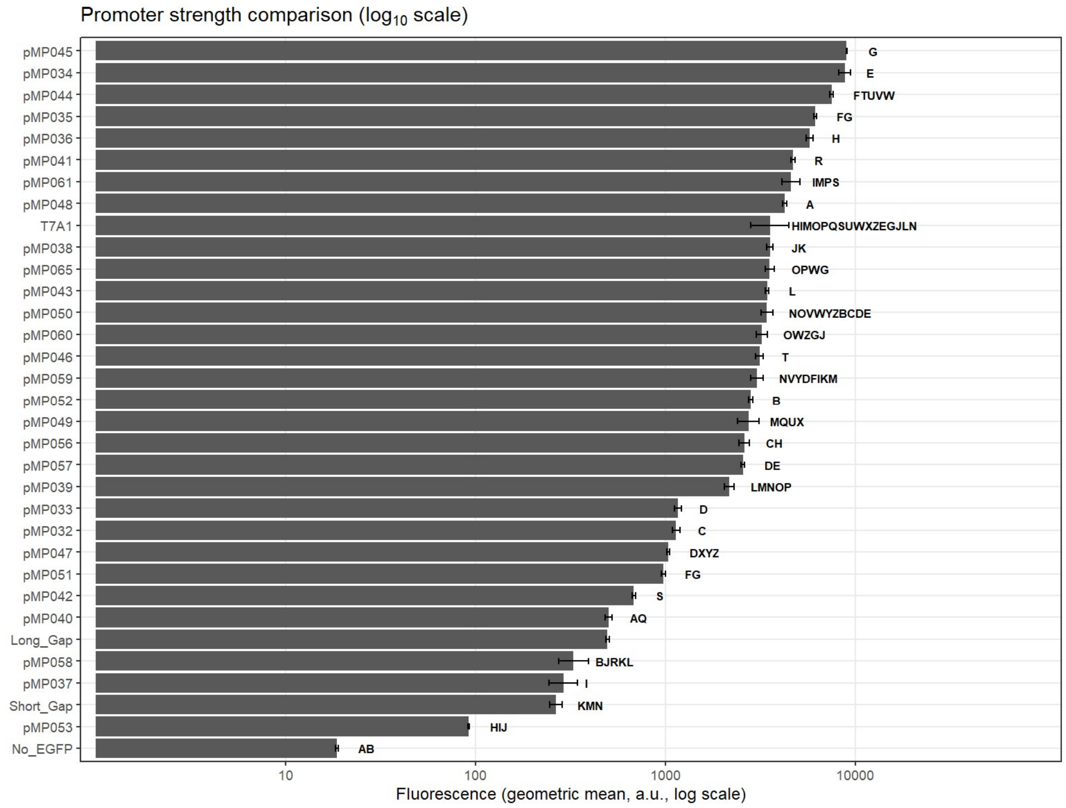


**Figure S4. Expression groups following post-hoc Welch pairwise tests with Benjamini–Hochberg correction**. Bars represent geometric mean fluorescence (± geometric SD; n = 3). Values were analysed on $\log_{10}$-transformed data by Welch's ANOVA with Benjamini–Hochberg–adjusted post-hoc tests. Compact letter displays (CLD) indicate significance groups (same letter = not significantly different, $p \geq 0.05$).

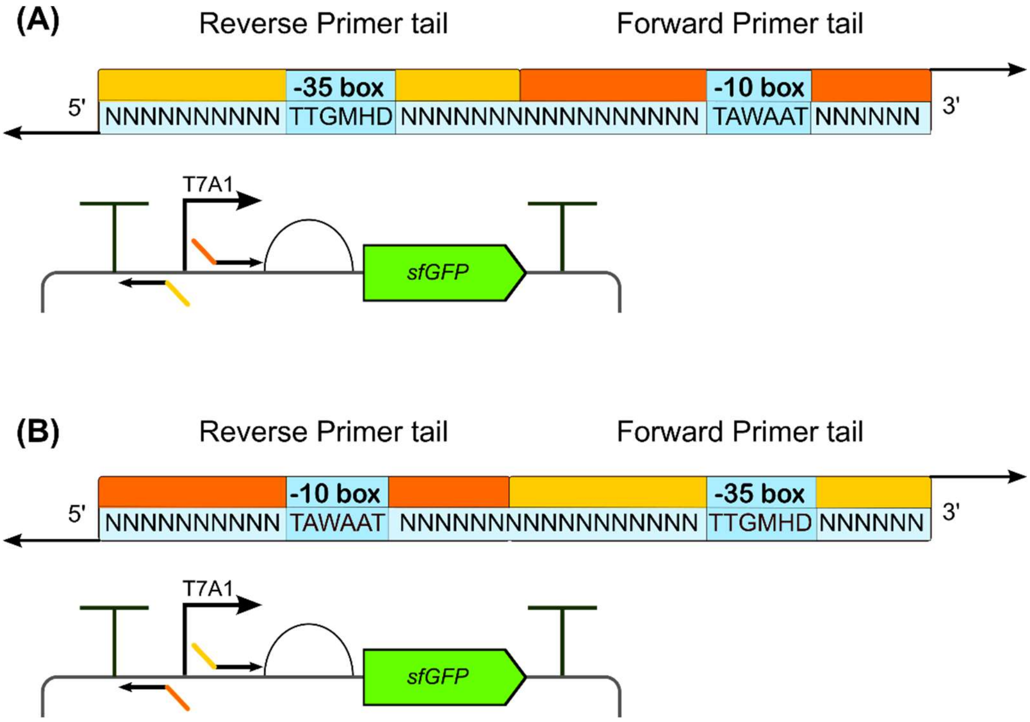


**Figure S5. (A) Planned synthetic promoter design.** The reverse primer was supposed to carry the -35 box, and the forward primer the -10 box. **(B) Actual synthetic promoter design.** The -10 box was carried by the reverse primer, and the -35 by the forward primer, leading to the swap of the operator sequences.

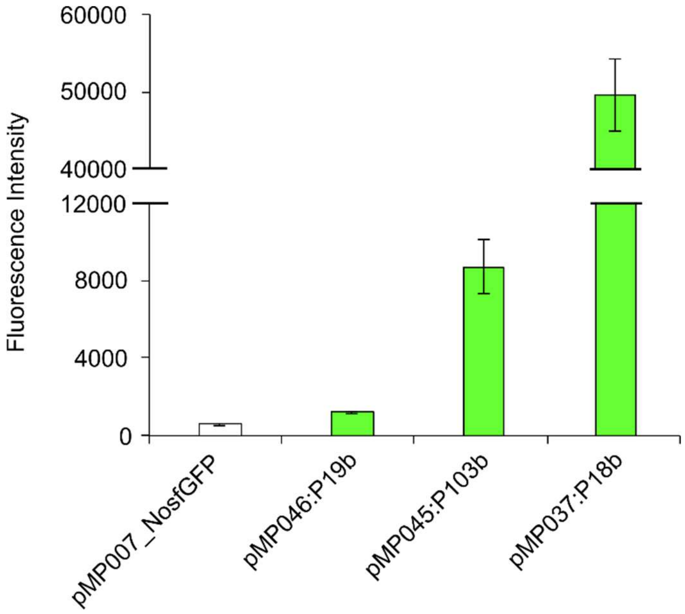


**Figure S6. Fluorescence intensity of sfGFP under promoters with standard order of operators.** Plasmids pMP046:P19b, pMP045:P103b and pMP037:P18b were tested by flow-cytometry for sfGFP fluorescence intensity along the negative control pMP007_No*sfGFP*. The plasmids pMP046:P19b, pMP045:P103b and pMP037:P18b respectively carried the versions of the weak, medium and strong promoters P19, P103 and P18 with the standard order of –35 and –10 boxes. *Z. mobilis* recombinant strains were grown aerobically in ZRMG rich medium, at 30°C, in 96-well plates, shaking at 800 rpm. Samples were harvested during late exponential phase. Error bars represent the standard deviation of three independent biological replicates.

**Table S3. Intrinsic terminators that were screened in *Z. mobilis*.**

| Terminator | Plasmid | Sequence (5’-3’) | Source | Δ*G* (kcal/mol) | Reference |
|---|---|---|---|---|---|
| ZM010 | pMM007 | GAAGGCGATGTGCTTTTGAGCATTATCCCTGT | *Z. mobilis* ZM4 | -10.7 | This study |
| ZM032 | pMM008 | GCAAGGCATCCGGTTTTAAATCTGGTAAATTG | *Z. mobilis* ZM4 | -6.6 | This study |
| ZM048 | pMM009 | AGACGGTGATCATGAAAAAAATTCTGCTTTTA | *Z. mobilis* ZM4 | -7.6 | This study |
| ZM050 | pMM010 | AGTTTAGGTAAAACGCCTTTGGCCGGCATTTA | *Z. mobilis* ZM4 | -5.7 | This study |
| ZM063 | pMM011 | AGTTGGTTTTTTTCGAGCTTTGAGTTTGGGAA | *Z. mobilis* ZM4 | -10.3 | This study |
| ZM065 | pMM012 | GCTGGAAACCTATCGTAACGCTTTGGGCTTGA | *Z. mobilis* ZM4 | -11 | This study |
| ZM068 | pMM013 | GCCTTTGAGTCGAGCCTTTGAGTCGAGCCTTT | *Z. mobilis* ZM4 | -8.6 | This study |
| L3S2P51 | pMM015 | CTCGGTACCAAAAAAAAAAAAAAAAGACGCTG | Synthetic | -24.9 | [44] |
| L3S1P56 | pMM016 | TTTTCGAAAAAAGGCCTCCCAAATCGGGGGG | Synthetic | -28.8 | [44] |
| ECK120033737 | pMM017 | GGAAACACAGAAAAAAGCCCGCACCTGACAG | *E. coli* K12 | -25 | [44] |

| | | | | | |
|---|---|---|---|---|---|
| ECK120015170 | pMM018 | ACAATTTTCGAAAAAACCCGCTTCGGCG GGTT | *E. coli* K12 | -20.1 | [44] |
| ECK120017009 | pMM019 | GATCTAACTAAAAAGGCCGCTCTGCGGC CTTTTTTCTTTTCACT | *E. coli* K12 | -16.2 | [44] |
| ECK120035133 | pMM020 | ACTGATTTTTAAGGCGACTGATGAGTCG CCTTTTTTTTGTCT | *E. coli* K12 | -15.4 | [44] |
| ECK120029600 | pMM021 | TTCAGCCAAAAAACTTAAGACCGCCGGT CTTGTCCACTACCTTGCAGTAATGCGGT GGACAGGATCGGCGGTTTTCTTTTCTCT TCTCAA | *E. coli* K12 | -42 | [44] |
| ECK120033736 | pMM022 | AACGCATGAGAAAGCCCCCGGAAGATCA CCTTCCGGGGGCTTTTTTATTGCGC | *E. coli* K12 | -37.8 | [44] |
| ECK120034435 | pMM023 | CTCGGTACCAAATTCCAGAAAAGAGACG CTGAAAAGCGTCTTTTTTCGTTTTGGTCC | *E. coli* K12 | -27.9 | [44] |

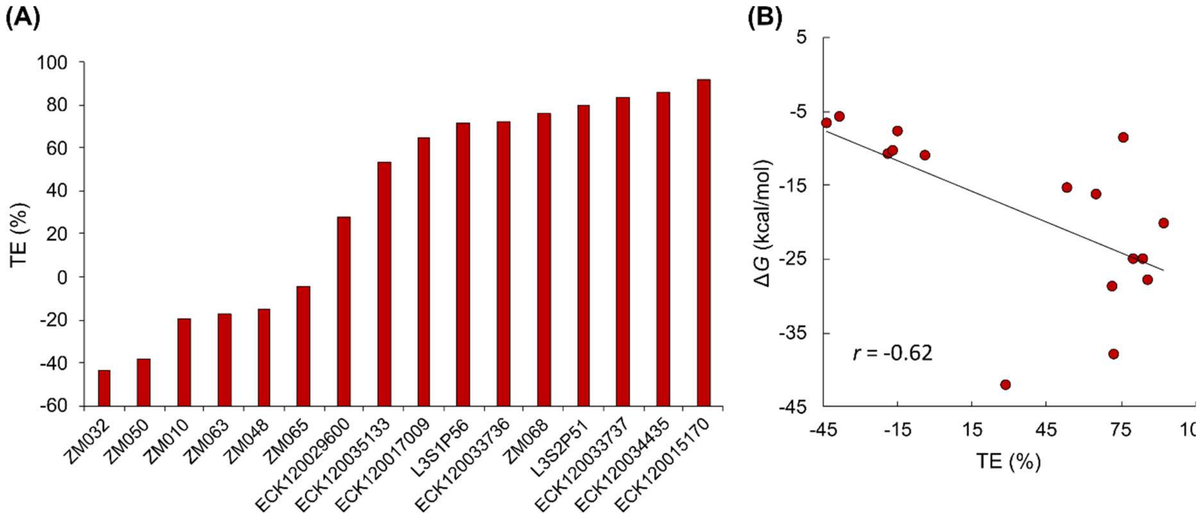


**Figure S7 Termination efficiency and their correlation to Δ*G*. (A)** Termination efficiency (TE) expressed in percentage of putative native terminators and synthetic and heterologous (*E. coli*) terminators in *Z. mobilis*. **(B) Correlation of Δ*G* and TE.** The scatter plot displays a moderate inverse relationship between Δ*G* and TE (Pearson correlation coefficient *r* = -0.62.

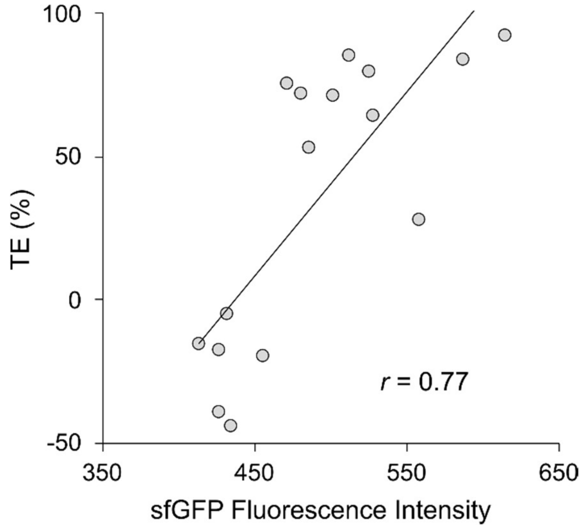


**Figure S8 Correlation of TE and sfGFP fluorescence intensity.** The fluorescence of the sfGFP increased with the strength of the terminator (Pearson correlation coefficient *r* = -0.77).

**Table S4. Spacers used in this study.**

| Spacer | Sequence (5’-3’) |
|---|---|
| **2** | CGCCCCCGGAGGCTTTCCCGGGGCAAATCA |
| **4** | ATCTCAACTCTTTTAAACAAGCGCGAGTCA |
| **5** | GTTGGTTTTTTTCGAGCTTTGAGTTTGGGAAA |
| **6** | ATTTGTAAAGTGTCTATTATCCCTAAGCCCATTTTTTGCA |

**Table S5. BCD sequences used in this study.**

| BCD | Sequence (5'-3') |
|---|---|
| BCD22 | GGGCCCAAGTTCACTTAAAAAGGAGATCAACAATGAAAGCAATTTTCGTACTGAAACATCTTAATCATGCCTAGGAAGTTTTCTAATG |
| BCD12 | GGGCCCAAGTTCACTTAAAAAGGAGATCAACAATGAAAGCAATTTTCGTACTGAAACATCTTAATCATGCTGCGGAGGGTTTCTAATG |
| BCD2 | GGGCCCAAGTTCACTTAAAAAGGAGATCAACAATGAAAGCAATTTTCGTACTGAAACATCTTAATCATGCTAAGGAGGTTTTCTAATG |

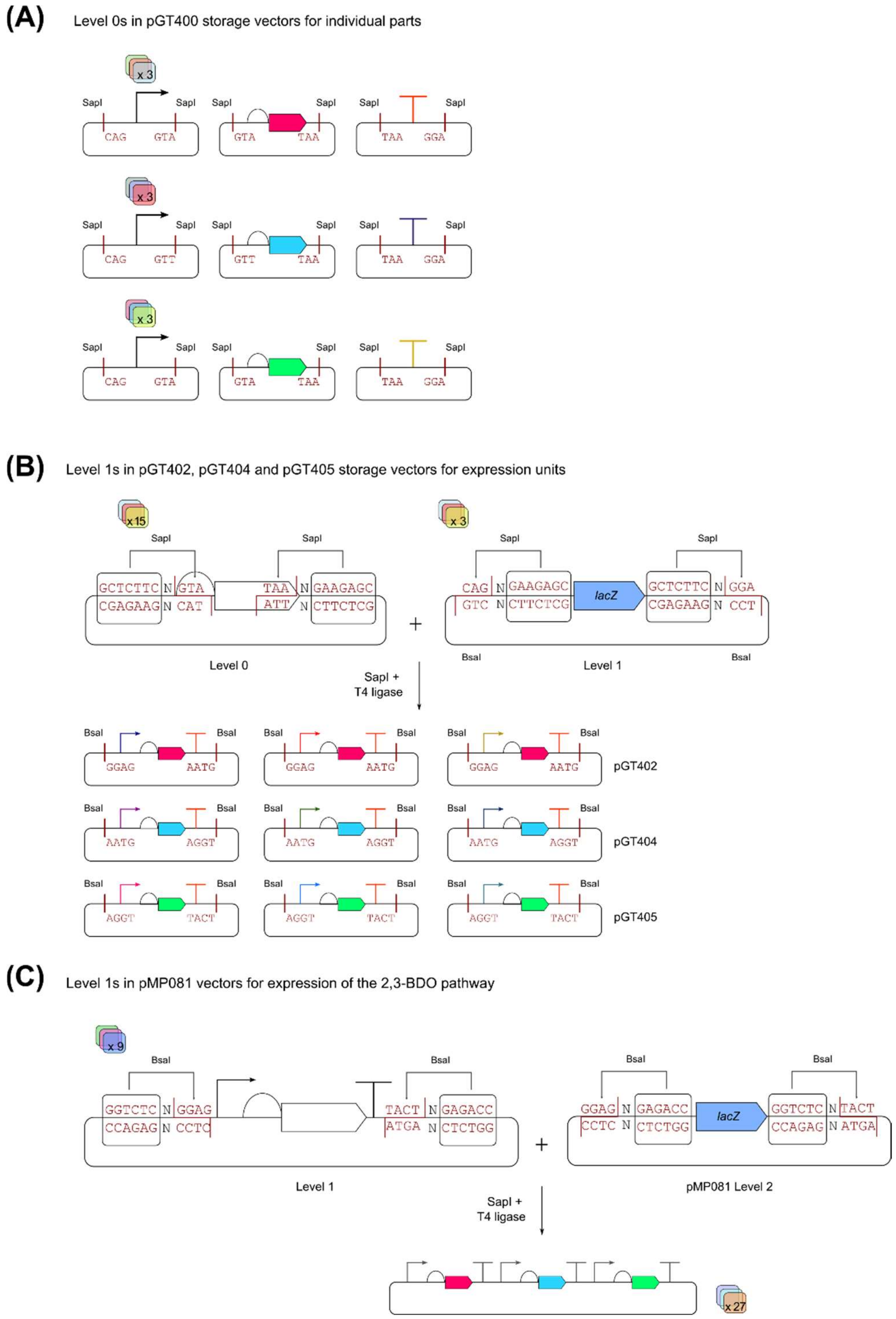


**Figure S9. Start-Stop assembly for building differentially regulated pathways. (A)** nine different promoters, three genes with their RBSs, and three strong terminators were cloned in the Level 0 pGT400. The inserts are flanked by inward-facing SapI restriction sites. **(B)** Promoters, genes and terminators were assembled into expression units in Level 1 plasmids (pGT402, pGT404 and pGT405), creating three different versions of transcriptionally regulated genes. Prior to digestion and ligation, the Level 1 plasmids carry a *lacZ* gene flanked by two-outward facing SapI sites. The fusion sites in the Level 1 plasmids are complementary to those carried by the start and end of the expression unit (promoter and terminator, respectively). Also, Level 1 plasmids have two inward-facing BsaI sites, which were used for the final assembly of the expression units into a multi-gene pathway. (C) The nine combinations of Level 1 plasmids were BsaI-cut and ligated in the Level 2 vector pMP081, which has outward-facing BsaI sites that create fusion sites for the binding of the first and last expression unit, generating up to twenty-seven differentially regulated pathways.

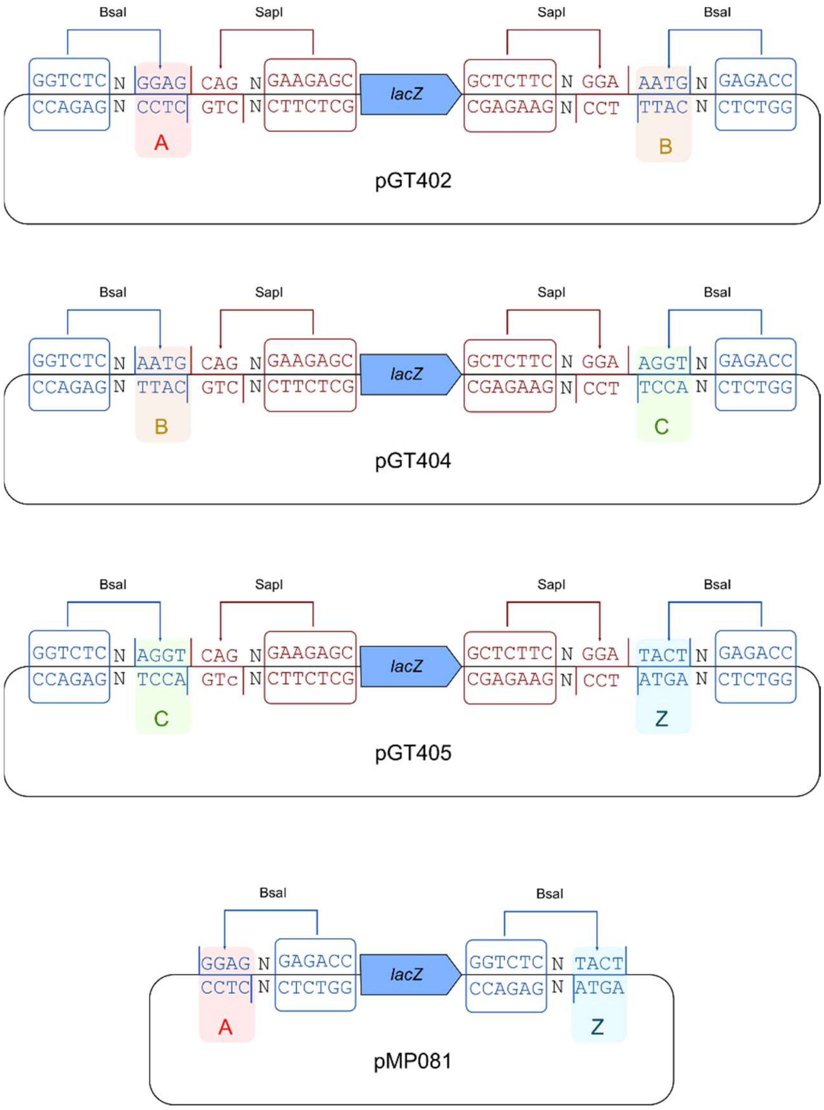


**Figure S10. SapI and BsaI restriction sites architecture in the Level 1 plasmids pGT402, pGT404 and pGT405, and in the final Level 2 vector pMP081.** The Level 1 plasmids carry two outward-facing SapI sites for the introduction of the expression units carrying complementary tails. Outside the SapI sites, two inward-facing BsaI sites generated the fusion site A and B, B and C, C and Z, for the sequential ligation of the expression units into the acceptor vector pMP081, which possesses the A and Z fusion sites.

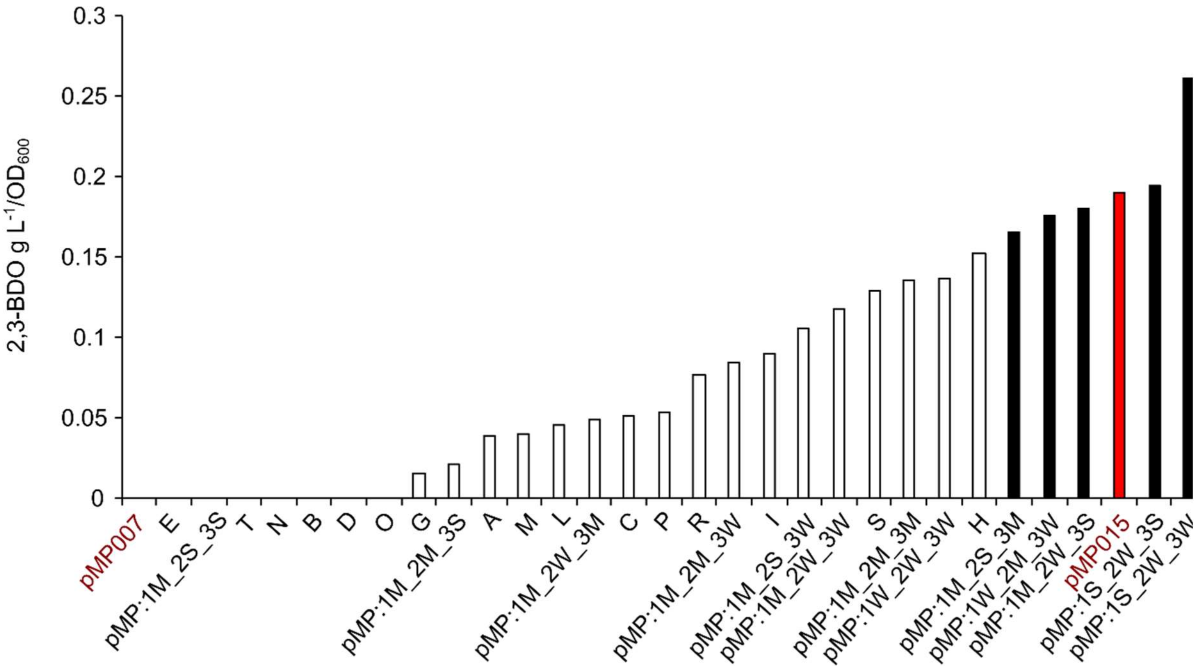


**Figure S11. Bar charts showing 2,3-BDO production in *Z. mobilis* strains after six days of growth.** Strains included the negative control (pMP007), reference control (pMP015), and thirty recombinant strains carrying plasmids for 2,3-BDO pathway expression. Cultures were grown in ZMMG80 minimal medium in 50 ml Falcon tubes at 30 °C, shaking at 120 rpm. Plasmids labelled with single capital letters were unsequenced. Sequenced plasmids were renamed as pMP and annotated according to the promoter strength associated to each gene of the pathway: 1 = *aldC*, 2 = *bdh*, 3 = *alsS*; W = weak, M = medium, S = strong. The top five 2,3-BDO producers were (pMP:1S_2W_3W), (pMP:1S_2W_3S), (pMP:1M_2W_3S), (pMP:1W_2M_3W), and (pMP:1M_2S_3M), excluding the control pMP015, which ranked third. Data are shown normalized by $OD_{600}$.

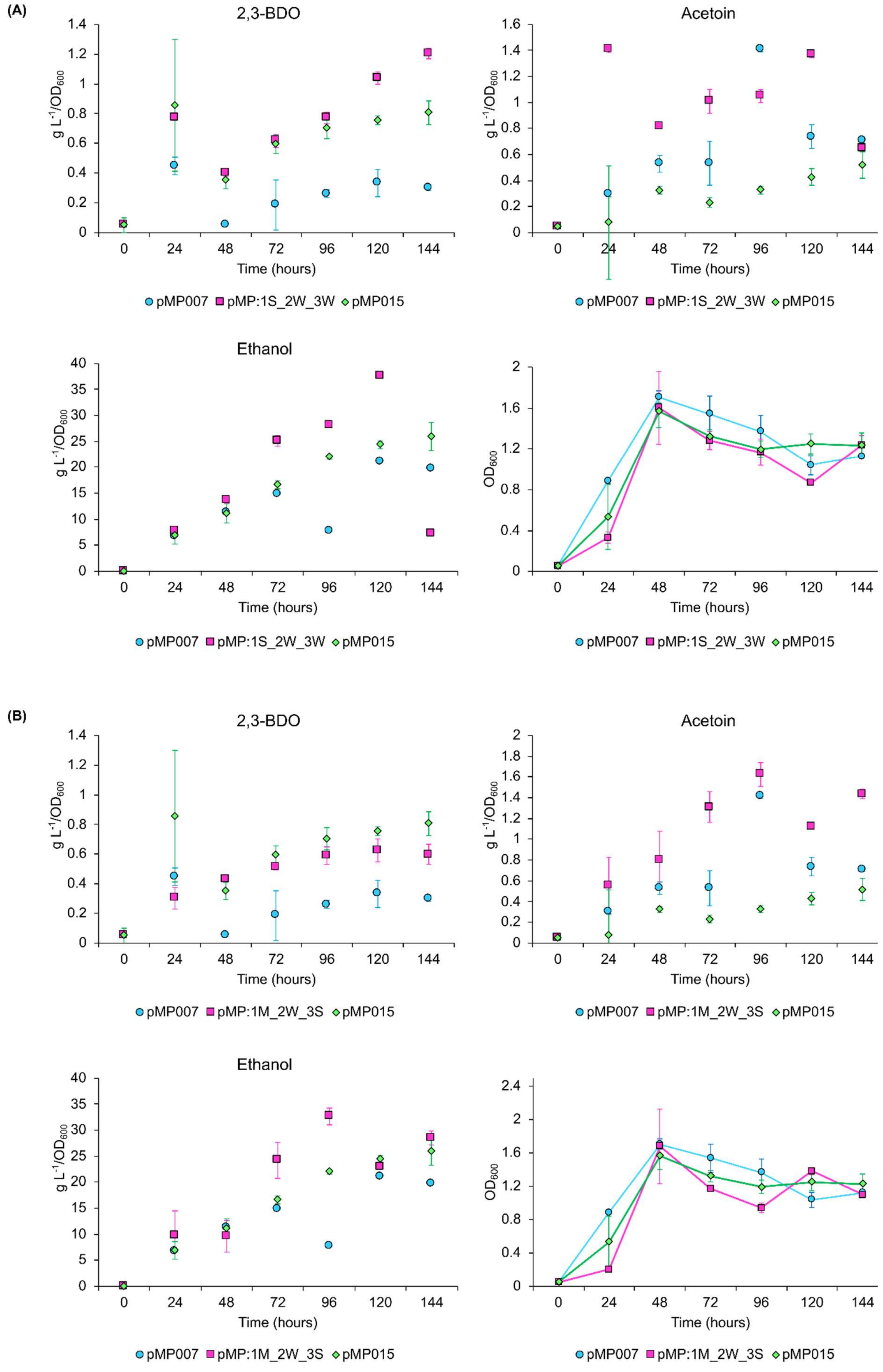
(A)
2,3-BDO
Acetoin
Ethanol
g L-1/OD600
OD600
Time (hours)
pMP007
pMP:1S_2W_3W
pMP015
(B)
2,3-BDO
Acetoin
Ethanol
g L-1/OD600
OD600
Time (hours)
pMP007
pMP:1M_2W_3S
pMP015

(C)

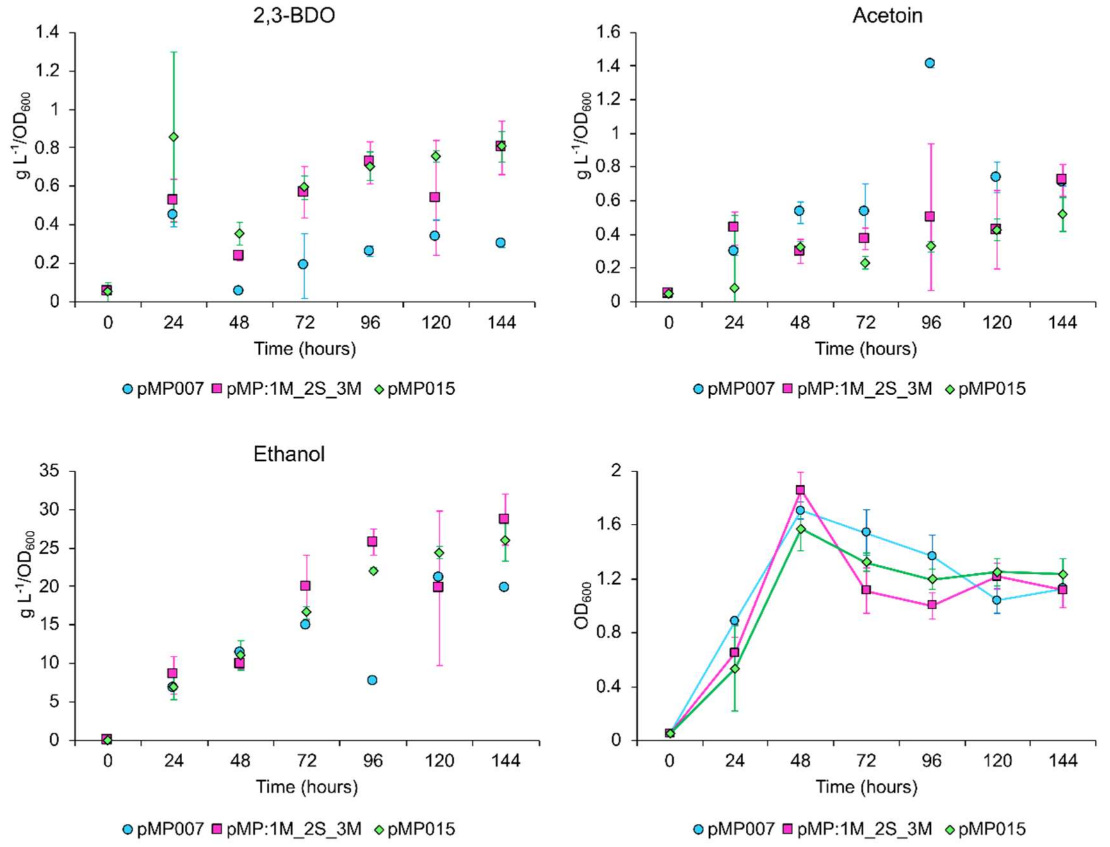

2,3-BDO
Acetoin
Ethanol
g L-1/OD600
OD600
Time (hours)
pMP007 pMP:1M_2S_3M pMP015


(D)

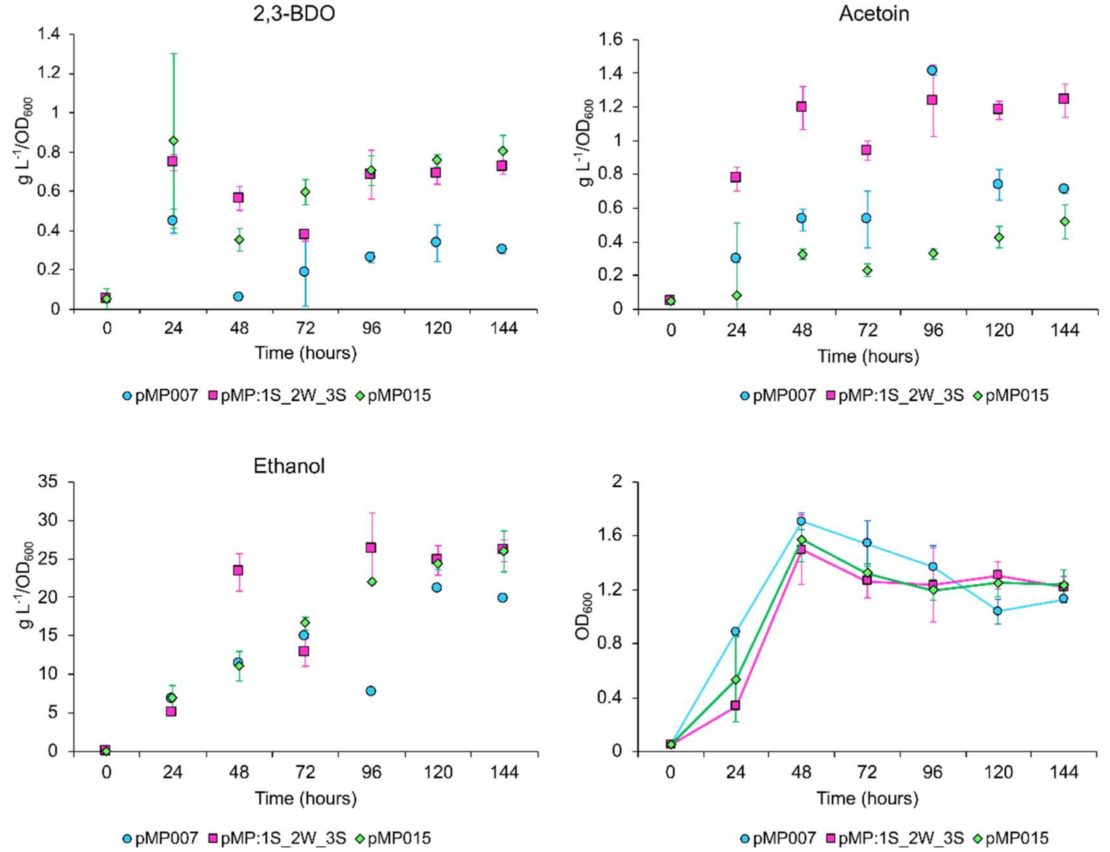

2,3-BDO
Acetoin
Ethanol
g L-1/OD600
OD600
Time (hours)
pMP007 pMP:1S_2W_3S pMP015

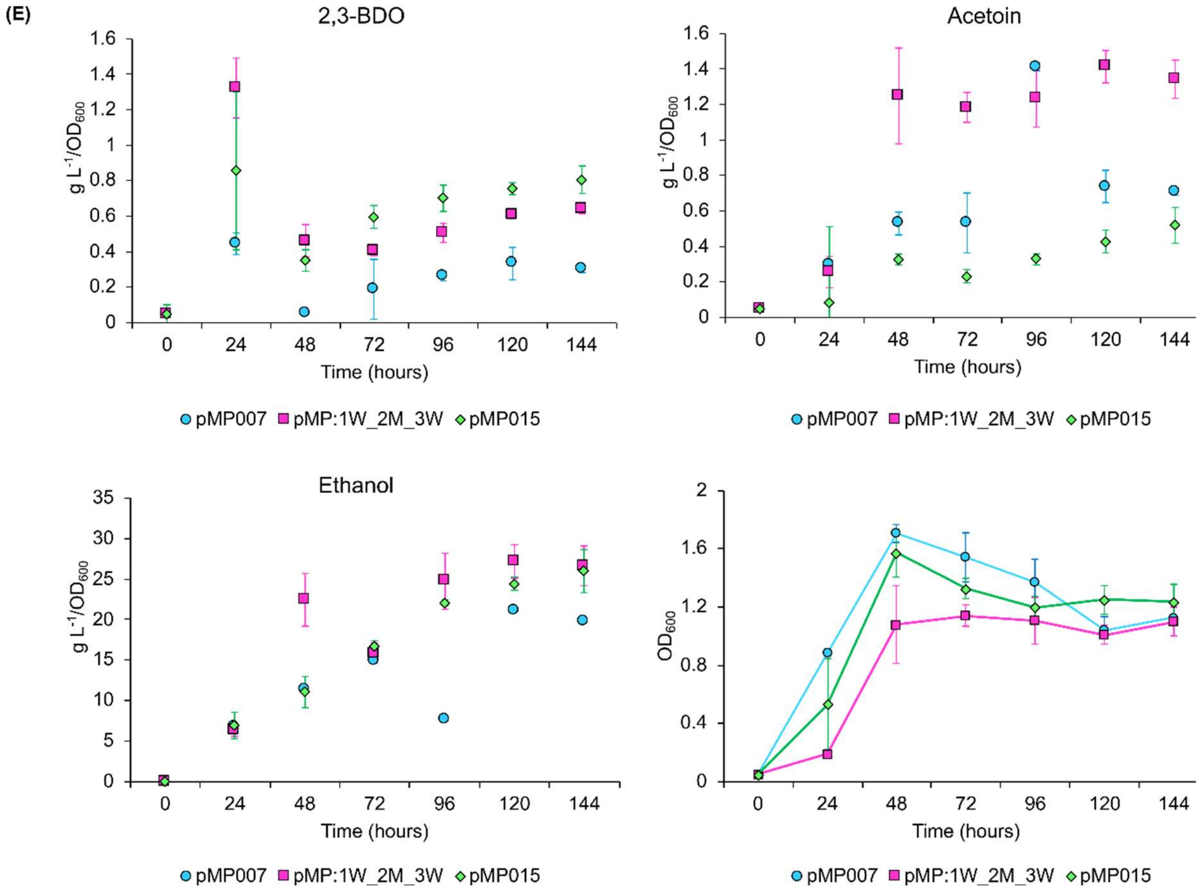


**Figure S12. 2,3-BDO, acetoin, ethanol production and growth of five *Z. mobilis* recombinant strain with differentially regulated 2,3-BDO pathways, negative control *Z. mobilis* (pMP007) and positive control *Z. mobilis* (pMP015). (A), (B), (C), (D)** and **(E)** panels respectively show 2,3-BDO, acetoin, ethanol production and growth of *Z. mobilis* (pMP:1S_2W_3W), (pMP:1M_2W_3S), (pMP:1M_2S_3M), (pMP:1S_2W_3S) and (pMP:1W_2M_3W) in pink squares, compared to the negative control (pMP007) in blue circles, and the positive control (pMP015) in green diamonds. 2,3-BDO, acetoin and ethanol were measured by HPLC, and concentrations were normalised by $OD_{600}$. Error bars represent the standard deviation of three independent biological replicates. Cultures were grown in ZMMG80, shaking at 120 rpm over 6 days (144 h).

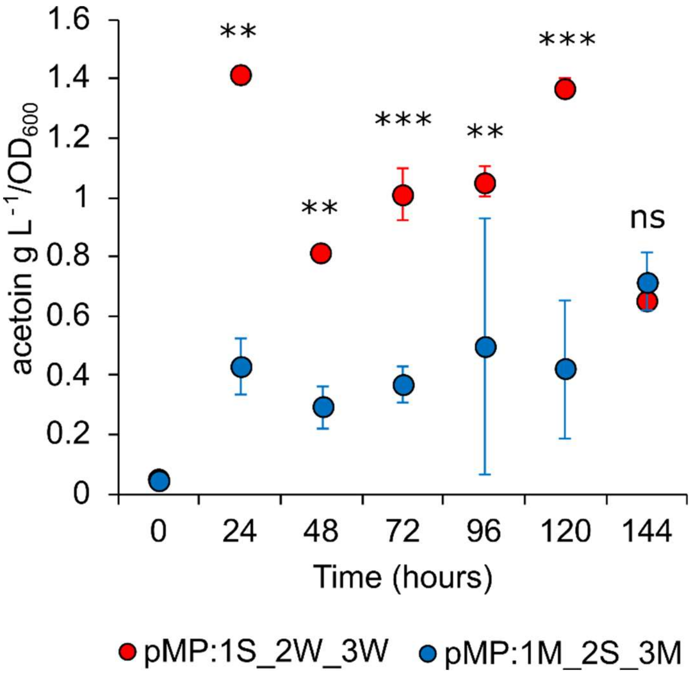


**Figure S13. Comparison of acetoin production between *Z. mobilis* (pMP:1S_2W_3W), in red, and (pMP:1M_2S_3M), in blue, over six days of growth in ZMMG80, shaking at 120 rpm.** Acetoin was measured by HPLC, and concentrations were normalised by $OD_{600}$. Error bars represent the standard deviation of three independent biological replicates. *** = $p < 0.01$, ** = $p < 0.05$, ns = not significant, two-sample *t*-test.

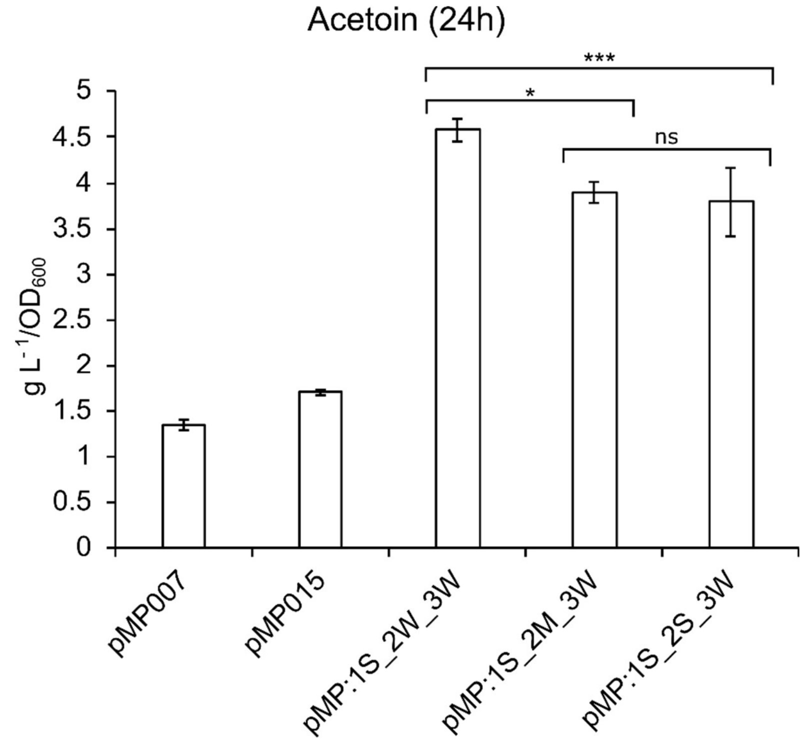


**Figure S14. Acetoin production in *Z. mobilis* negative (pMP007) and positive (pMP015) controls, and recombinant strains (pMP:1S_2W_3W), (pMP:1S_2M_3W), and (pMP:1S_2S_3W).** Cultures were grown in ZRMG rich medium, and samples were collected after 24 hours. Acetoin concentrations were determined by NMR and normalized to OD600. Error bars represent the standard deviation of three biological replicates. Statistical significance was assessed using two-sample *t*-tests: pMP:1S_2W_3W vs pMP:1S_2M_3W *p* value = 0.04, pMP:1S_2W_3W vs pMP:1S_2S_3W *p* value = 0.005, and pMP:1S_2M_3W vs pMP:1S_2S_3W *p* value= 0.71.
